# Rotavirus Induces Intercellular Calcium Waves through ADP Signaling

**DOI:** 10.1101/2019.12.31.892018

**Authors:** Alexandra L. Chang-Graham, Jacob L. Perry, Melinda A. Engevik, Heather A. Danhof, Francesca J. Scribano, Kristen A. Engevik, Joel C. Nelson, Joseph S. Kellen, Alicia C. Strtak, Narayan P. Sastri, Mary K. Estes, James Versalovic, Robert A. Britton, Joseph M. Hyser

**Affiliations:** Department of Molecular Virology and Microbiology, Baylor College of Medicine; Department of Pathology and Immunology, Baylor College of Medicine; Department of Pathology, Texas Children’s Hospital; Department of Medicine, Gastroenterology and Hepatology, Baylor College of Medicine

## Abstract

Rotavirus causes severe diarrheal disease in children by broadly dysregulating intestinal homeostasis. However, the underlying mechanism(s) of rotavirus-induced dysregulation remains incompletely characterized. Here we show that rotavirus-infected cells produce paracrine signals that manifest as intercellular calcium waves (ICWs); which are observed in both cell lines and human intestinal enteroids. Rotavirus ICWs are caused by the release of extracellular adenosine diphosphate (ADP) that activates P2Y1 purinergic receptors on neighboring cells and are blocked by P2Y1 antagonists or CRISPR/Cas9 knockout of P2Y1. Blocking the paracrine ADP signal reduces rotavirus replication, inhibits rotavirus-induced serotonin release and fluid secretion, and reduces diarrhea severity in neonatal mice. This is the first evidence that viruses exploit ICWs to amplify diarrheal signaling; a finding which has broad implications for gastrointestinal physiology.

**Summary:** Rotavirus triggers the extracellular release of ADP from infected cells to dysregulate nearby uninfected cells and activate pro-disease pathways.

## Introduction

Rotavirus (RV), a non-enveloped double-stranded RNA virus of the *Reoviridae* family, causes severe diarrhea and vomiting in children worldwide, resulting in ∼258 million diarrhea episodes and ∼128,000 deaths annually (*1*). Though the pathways by which other enteric pathogens cause diarrhea are well-characterized, such as cholera toxin of *Vibrio cholerae*, the mechanisms of RV diarrhea are multifactorial and still not completely understood. RV infects the epithelial enterocytes and enteroendocrine cells at the tip and middle of villi in the small intestine (*2*–*4*). Notably, only a percentage of cells susceptible to RV are infected, and diarrhea occurs before the onset of histopathologic changes (*2, 5*–*7*). During infection, RV dysregulates host cell calcium (Ca^2+^) signaling pathways to increase cytosolic Ca^2+^, which is required for RV replication. The RV nonstructural protein 4 (NSP4) drives these changes in Ca^2+^ homeostasis as both an intracellular viral ion channel (*i.e.*, viroporin) in the endoplasmic reticulum and an extracellular enterotoxin that elicits a phospholipase-C-dependent increase in cytosolic Ca^2+^ (*8*–*10*). These perturbations to host Ca^2+^ signaling are known to activate autophagy, disrupt the cytoskeleton and tight junctions, and trigger fluid secretion pathways (*9, 11*–*14*).

A long-held concept in how RV infection causes life-threatening diarrhea, despite a small percentage of infected cells, is that RV-infected cells release potent signaling molecules that can dysregulate neighboring, uninfected cells (*6, 15, 16*). This notion is based on previous studies that have identified increases in candidate signaling molecules during RV infection, including enterotoxin NSP4, prostaglandins (PGE2), and nitric oxide (NO) (*9, 17*–*19*). In this theory, enterotoxin NSP4 can bind to neighboring, uninfected enterocytes to activate Ca^2+^-activated chloride channels and cause secretory diarrhea (*20*–*22*), and PGE2 and NO may further activate fluid secretion processes (*23, 24*). Simultaneously, dysregulation of neighboring enteroendocrine cells triggers the Ca^2+^-dependent release of serotonin, which stimulates the enteric nervous system both to activate vomiting centers in the central nervous system and to activate secretory reflex pathways in the gastrointestinal (GI) tract (*4, 7*). Thus, this concept of RV-induced diarrhea addresses how limited infection at the middle-to-upper villi may cause widespread dysregulation of host physiology and life-threatening disease. However, this cell-to-cell functional signaling has not been directly observed in a RV infection, and the signaling pathways remain incompletely understood.

Herein we demonstrate that RV-infected cells signal to uninfected cells via an extracellular purinergic signaling pathway. This newly identified pathway is a dominant driver of observed RV disease processes, including replication, upregulation of PGE2- and NO-producing enzymes, serotonin secretion, fluid secretion, and diarrhea in a neonatal mouse model. Our findings provide new insights into the mechanism(s) of viral diarrhea and gastrointestinal physiology.

## Results

### Low multiplicity infection reveals intercellular calcium waves

Previous studies have shown that RV significantly increases cytosolic Ca^2+^ during infection and disrupts host Ca^2+^-dependent processes to cause disease (*25*–*27*). We used African Green monkey kidney MA104 cells stably expressing the genetically-encoded calcium indicator (GECI) GCaMP5G or GCaMP6s to observe changes in cytosolic Ca^2+^ during RV infection using live-cell time-lapse epifluorescence imaging. We mock- or RV (strain SA114F)-infected MA104-GCaMP cells at a low multiplicity of infection (MOI 0.01) to distinguish Ca^2+^ changes in infected cells compared to neighboring, uninfected cells. RV-infected cells were identified using immunofluorescence (IF) staining for RV antigen after imaging, and these IF images were superimposed on the time-lapse movies (**Fig. 1A**). While mock-infected cells had minimal changes in GCaMP fluorescence, RV-infected cells exhibited large fluctuations in fluorescence, and thus increases in cytosolic Ca^2+^, as discrete signaling events, consistent with previous findings (*28*). These signaling events not only increased cytosolic Ca^2+^ in the RV antigen-positive cell, but we also observed successive increases in cytosolic Ca^2+^ in neighboring, uninfected cells over a time interval of 30 s (**Fig. 1A, Supplementary Movie S1**). We quantified the Ca^2+^-eliciting signal from infected to uninfected cells using a previously described criterion in which changes in normalized relative fluorescence greater than 5% constituted a “Ca^2+^ spike” (*28*). We observed that with neighboring cells 5 (NB+5) and 10 (NB+10) cells away from the infected cell, the number and average magnitude of the RV-induced Ca^2+^ transients decreased with increasing distance from the infected cell (**Fig. 1B, blue and purple traces**). Quantification of the number and magnitude of Ca^2+^ spikes of the infected, the NB+5, and the NB+10 cells show a significant decrease with distance from the infected cell (**Fig. 1C**,**D**). Together, this data constitutes the first direct observation of a RV-infected cell signaling to surrounding uninfected cells and eliciting a Ca^2+^ signaling response.

**Figure 1.**
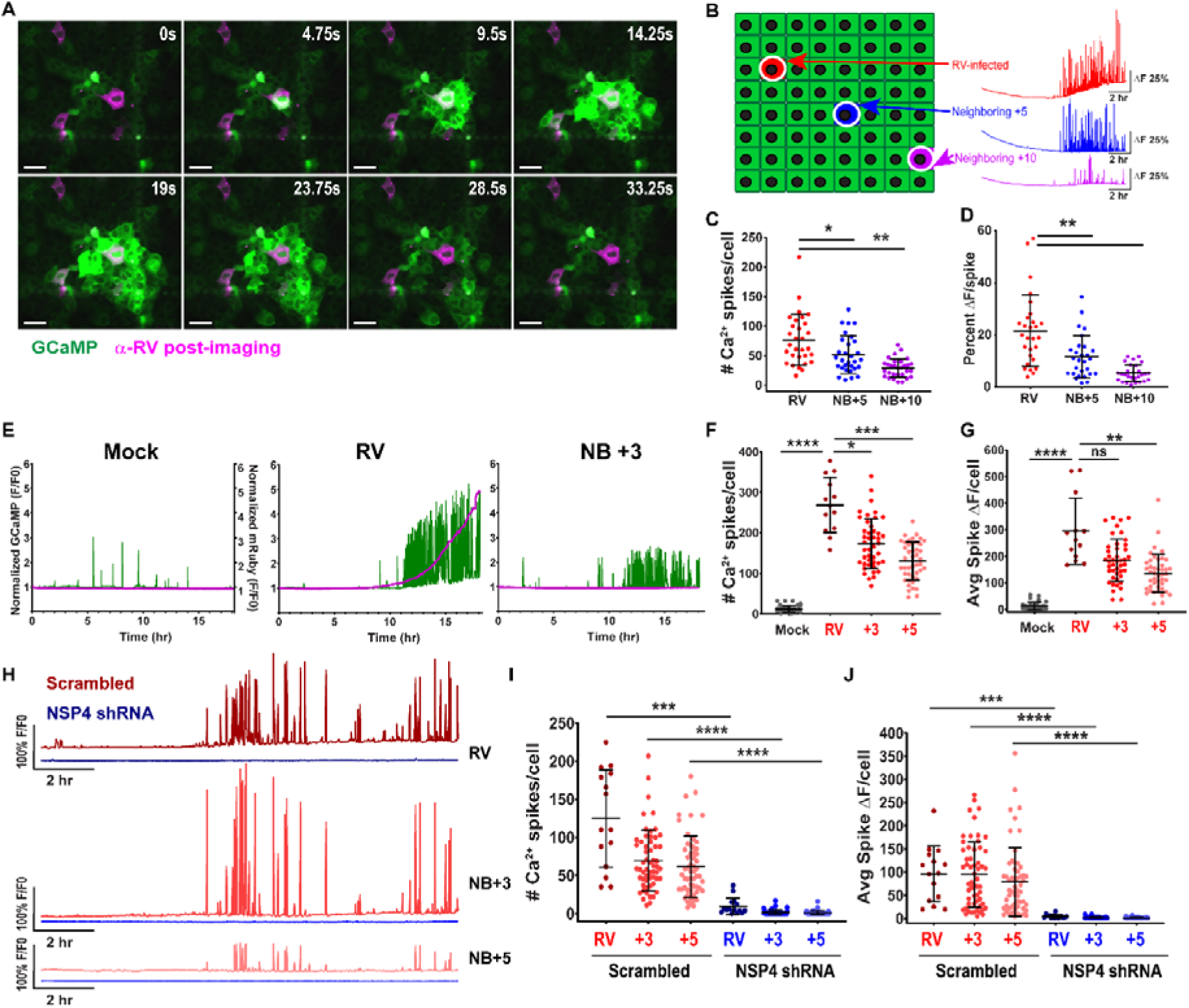
Rotavirus-infected cells trigger calcium signaling in neighboring uninfected cells. **(A)** MA104-GCaMP cells, RV (SA114F)-infected (pink). Scale bar = 100 µm. **(B)** Representative Ca^2+^ traces of normalized GFP fluorescence (F/F_0_) of RV-infected cell and neighboring (NB) cells. **(C, F, I)** Ca^2+^ spikes per cell and **(D, G, J)** average F/F_0_ increase of Ca^2+^ spike/cell **(E-G)** Mock- or RV (SA11cl3-mRuby3)-infected, **(E)** representative Ca^2+^ traces Left axis: GCaMP F/F_0_ (green), Right axis: mRuby3 F/F_0_ (pink). **(H-J)** MA104-GCaMP-shRNA scrambled or shRNA-NSP4 cells infected with RV (SA11cl3-mRuby3), **(H)** Representative Ca^2+^ traces. Kruskal-Wallis with Dunn’s multiple comparisons test. (*p<0.05, **p<0.01, ***p<0.001, ****p<0.0001)

Next, we sought to confirm that the Ca^2+^ signaling in the neighboring cells was not due to RV infection undetected by our antibody staining. MA104-GCaMP cells were mock-inoculated or infected with a recombinant SA11 clone 3 strain containing an mRuby3 fluorescent reporter (SA11cl3-mRuby3) as described previously (*28*), at MOI 0.01 and imaged for ∼3-22 hours post-infection (hpi), conditions we used throughout these studies unless noted otherwise. We found no increase in mRuby3 fluorescence in the mock-infected cells but a steep increase in fluorescence in RV-infected cells by hour 12 of imaging that correlated with increased Ca^2+^ signaling (**Fig. 1E, left and middle panels, Supplementary Movie S2**). Importantly, the NB+3 cell also had robust Ca^2+^ signaling by hour 12, but the lack of increase in mRuby3 fluorescence demonstrated it was not infected (**Fig. 1E, right panel**). As before, there was a similar decrease in the number of Ca^2+^ spikes and average Ca^2+^ spike magnitude in the NB+3 and NB+5 cells that correlated with increasing distance from the infected cell (**Fig. 1 F, G**). These findings demonstrate that the Ca^2+^ signaling in the neighboring cell is not due to infection of that cell but is due to signals activated by a nearby RV-infected cell.

Since RV-induced changes in Ca^2+^ homeostasis are driven by NSP4, we next determined whether knockdown of NSP4 would reduce the Ca^2+^ signals in infected and uninfected cells. Using RV (SA11cl3-mRuby3), we infected MA104-GCaMP-shRNA cells expressing NSP4-targeted shRNA or a non-targeted scrambled shRNA described previously (*28*). RV infection of the MA104-GCaMP-shRNA cells with SA114F, which has the same NSP4 sequence as SA11cl3-mRuby3) decreases NSP4 expression by ∼85% and significantly decrease Ca^2+^ signaling (*28*). Here we observed that the Ca^2+^ signaling frequency and Ca^2+^ spike magnitudes were significantly decreased in the NSP4-targeted shRNA MA104-GCaMP cells, both in RV-infected cells and NB+3 and NB+5 uninfected cells (**Fig. 1H-J**). These results support that the RV-induced Ca^2+^ signal requires the expression of RV NSP4 and the dysregulation of host Ca^2+^ homeostasis in the infected cell.

### ADP and the P2Y1 receptor mediate RV-induced intercellular calcium waves

The pattern of signaling observed between RV-infected cells to uninfected cells is a phenomenon called intercellular calcium waves (ICWs), in which increases in cytosolic Ca^2+^ occur in an expanding circular pattern from a central initiating cell (*29*–*31*). Several molecules have been implicated in ICW propagation and may communicate via gap junctions or through paracrine signaling (*31*). To test if the RV-induced ICWs propagated via gap junctions, we infected MA104-GCaMP cells and treated with DMSO vehicle or the gap junction blockers 18β-glycyrrhetinic acid and TAT-Gap19. There was no significant difference in the Ca^2+^ spikes in neighboring cells between the mock-, 18β-glycyrrhetinic acid-, or TAT-Gap19-treated cells, supporting that the RV-induced ICWs did not propagate *via* gap junctions (**Fig. 2A**). This finding was consistent with a negative dye-loading scratch assay (**Supplementary Fig. S1A**). Next we tested if the extracellular enterotoxin form of NSP4 may be the signaling molecule responsible for propagating the ICWs by treating RV-infected MA104-GCaMP cells with purified anti-NSP4 rabbit antisera (120-147) known to bind to the enterotoxin domain(*22*) or purified anti-NSP4 monoclonal antibody 622 that neutralizes eNSP4-mediated Ca^2+^ flux (**Fig. 2B**). Addition of purified M60, a non-neutralizing anti-VP7 monoclonal antibody, was used as a control (*32*). We did not observe a significant reduction in Ca^2+^ spikes in the neighboring cells treated with anti-NSP4 antibodies, suggesting that enterotoxin NSP4 was not responsible for the RV-induced ICWs (**Fig. 2B, Supplementary Movie S3**). Previous studies found increased PGE2 and NO during RV infection, and thus are potential mediators of the RV-induced ICWs (*17*–*19*). However, adding PGE2 or the NO donor NOC-7 to MA104-GCaMP cells did not elicit a Ca^2+^ response, and thus these molecules cannot directly mediate the RV-induced ICWs (**Supplementary Fig. S1B-E**). Together these data support that ICW propagation is mediated by a signaling molecule not previously identified in RV infection.

**Figure 2.**
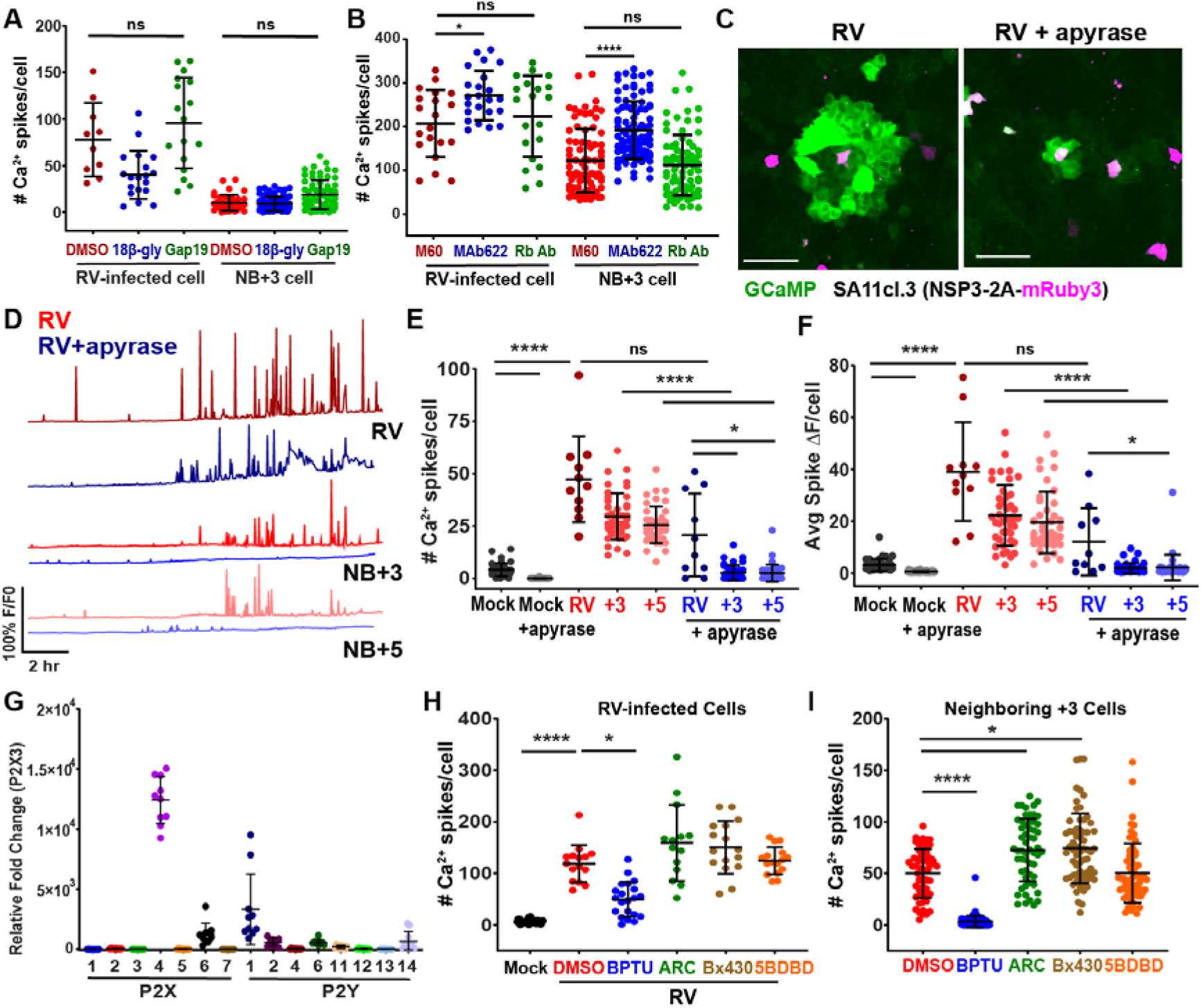
Rotavirus induces intercellular calcium waves by activating purinergic signaling. **(A-B)** Ca^2+^ spikes in MA104-GCaMP cells in RV-infected (SA11cl3-mRuby3) and neighboring (NB)+3 cell, treated with: **(A)** DMSO or gap junction blockers or **(B)** anti-VP7 M60 MAb, anti-NSP4 MAb 622, or anti-NSP4 antisera 120-147 (Rb Ab). **(C)** RV-infected cells, mock- or apyrase-treated and **(D)** representative Ca^2+^ traces. **(E)** Ca^2+^ spikes and **(F)** average F/F_0_ increase of Ca^2+^ spikes. **(G)** qPCR of purinergic receptor mRNA transcripts in MA104 cells. **(H-I)** Ca^2+^ spikes in RV-infected **(H)** or NB **(I)** cells treated with DMSO or purinergic receptor blockers. Kruskal-Wallis with Dunn’s multiple comparisons test. (Scale bar = 100 µm) (*p<0.05, ****p<0.0001)

We then sought to determine whether ATP, an important extracellular messenger for many cell types and a common mediator of ICW signaling, was the signaling molecule responsible for the RV-induced ICWs(*31*). We mock- or RV (SA11cl3-mRuby3)-infected MA104-GCaMP cells and mock-treated or treated with apyrase, an enzyme that degrades the purines ATP and ADP, after inoculation. Apyrase-treated wells showed a dramatic decrease in the area of Ca^2+^ signaling compared to untreated wells (**Fig. 2C, Supplementary Movie S4**). With apyrase treatment, the Ca^2+^ signaling frequency and amplitude are decreased both in the RV-infected cell and in neighboring, uninfected cells (**Fig. 2D**). The number of Ca^2+^ spikes and the average Ca^2+^ spike magnitude are significantly decreased in RV-infected wells treated with apyrase (**Fig. 2E-F**). These results support that extracellular ATP/ADP are the signaling molecules responsible for propagating ICWs in RV infection.

Purinergic signaling, *i.e.*, nucleotides as extracellular signals, is a key signaling pathway in intestinal physiology and involves multiple receptor subtypes. Using qPCR, we found that MA104 cells showed greatest expression of the purinergic receptors P2×4, P2×6, P2Y1, P2Y2, and P2Y14 (**Fig. 2G**). Relative mRNA expression levels of purinergic receptors did not substantially change with RV infection (**Supplementary Fig. S2A**). We first tested the ligand-gated ion channel P2×4 and the Gq/11 protein-coupled receptors P2Y1 and P2Y2 as they are strongly activated by ATP or ADP, which are well known to mediate ICWs (*31*). We treated RV(SA11cl3-mRuby3)-infected MA104-GCaMP cells with small molecule inhibitors of the receptors (Bx430 and 5BDBD for P2×4, BPTU for P2Y1, and AR-C 118925XX for P2Y2). The P2×4 inhibitors Bx430 and 5BDBD or the P2Y2 inhibitor AR-C 118925XX had no effect on RV-induced Ca^2+^ signaling. We found that the P2Y1 blocker BPTU significantly decreased Ca^2+^ spikes in both the RV-infected cell and in the NB+3 cells, and this decrease was dose-dependent (**Fig. 2H-I, Supplementary Fig. S2C, Supplementary Movie S5**). Additionally, we tested the specificity of the pharmacological blockers BPTU and AR-C 118925XX for their intended targets, the P2Y1 and P2Y2 receptors respectively, which are strongly activated by ADP and ATP, respectively. We observed that the P2Y1 inhibitor BPTU significantly reduced the Ca^2+^ response in MA104-GCaMP cells from ADP, but not when ATP was added (**Supplementary Fig. S2D**). Similarly, the P2Y2 inhibitor AR-C 118925XX significantly reduced the Ca^2+^ response from ATP, but not from ADP, supporting that these pharmacological inhibitors are specific for their targets (**Supplementary Fig. S2D**). Together these results support that the RV-induced ICWs are mediated by the extracellular release of ADP from infected cells and activation of the P2Y1 receptor on neighboring, uninfected cells.

### RV induces intercellular calcium waves in human intestinal enteroids

MA104 cells are a robust model system to study RV replication and form a simple flat epithelium that facilitates microscopy studies. However, they are neither human nor intestinal in origin. To determine whether RV also causes ICWs in intestinal epithelial cells, we used jejunum-derived human intestinal enteroids (HIEs) stably expressing the GECI GCaMP6s (jHIE-GCaMP6s) (*28*). HIEs are an *in vitro* model culture system established from human intestinal epithelial stem cells, making them a non-cancerous, non-transformed *in vitro* model of small intestine epithelium (*33*). Furthermore, HIEs recapitulate the different epithelial cell types and support RV infection and replication, making them a biologically relevant system to study the gastrointestinal epithelium and disease processes (*34*–*36*). We infected jHIE-GCaMP6s monolayers with mock inoculum or the human RV strain Ito and imaged at 4 hpi (**Fig. 3A, Supplementary Movie S6**). We observed ICWs in both mock- and RV-infected jHIE-GCaMP6s monolayers; however, the areas of Ca^2+^ signaling were typically larger and had greater frequency in the RV-infected monolayer. To verify infection, we fixed the monolayers at 24 hpi, immunostained for RV antigen, and observed positive staining in the RV-infected monolayers (**Fig. 3B**). These observations support that ICWs occur under basal conditions in human intestinal epithelial cell models, but infection with RV increases the size of the ICWs.

**Figure 3.**
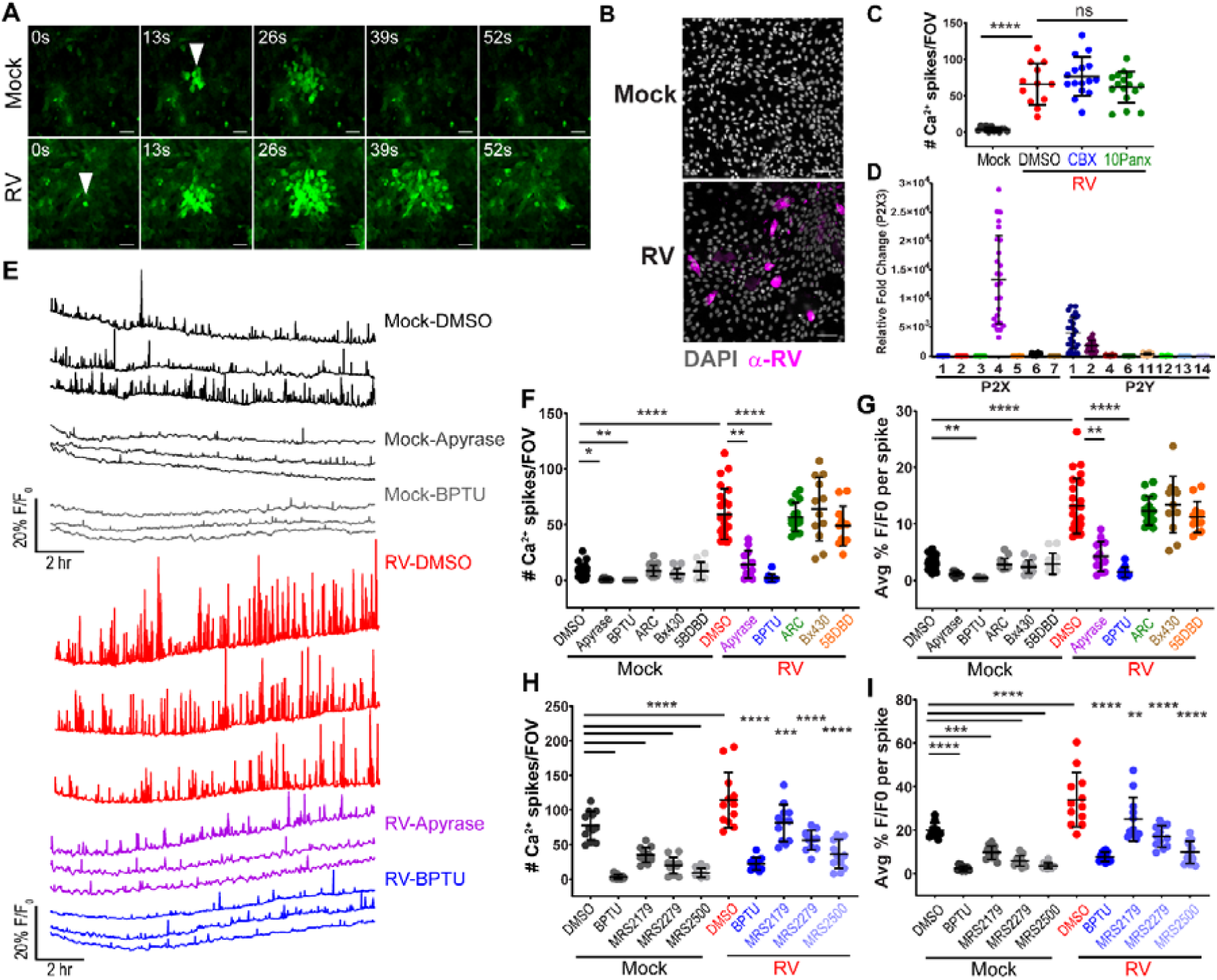
Rotavirus infection induces intercellular waves in human intestinal enteroids. **(A)** J3 jHIE-GCaMP6s monolayers mock- or RV(Ito)-infected (ICW, arrowhead) and **(B)** immunofluorescence. **(C)** Ca^2+^ spikes per field-of-view (FOV), in (J3)HIEs mock- or RV-infected and treated with vehicle, 100 µM carbenoxolone, or 10 µM 10Panx. **(D)** qPCR of purinergic receptor mRNA transcripts in (J3)HIEs. **(E-G)** Mock- or RV-infected (J3)HIEs, treated with DMSO, apyrase, or purinergic receptor blockers, **(E)** Ca^2+^ traces, **(F)** Ca^2+^ spikes/FOV, and **(G)** average F/F_0_ increase of Ca^2+^ spikes. **(H)** Ca^2+^ spikes/FOV of J2 jHIE-GCaMP6s monolayers mock- or RV-infected and treated with DMSO or P2Y1 receptor blockers, and **(I)** average F/F_0_ increase of Ca^2+^ spikes. One-way ANOVA with Bonferroni’s **(C**,**H**,**I)** and Kruskal-Wallis with Dunn’s **(F**,**G)** multiple corrections test. (*p<0.05, **p<0.01, ***p<0.001, ****p<0.0001) scale bar = 50 µm.

Next, we investigated the mechanism of action of the RV-induced ICWs in the HIEs. To test gap junction transmission, we treated mock- or RV (Ito)-infected jHIE-GCaMP6s monolayers with DMSO vehicle, the glycyrrhetinic acid-derivative carbenoxolone (CBX), or the pannexin blocker ^10^Panx. Since individual jHIE cells are much smaller and more mobile than MA104 cells, we were unable to perform single-cell analysis for these experiments. Instead we measured the GCaMP6s fluorescence over the whole field-of-view (FOV) (∼455 µm^2^). While we observed statistically greater Ca^2+^ spikes in RV-infected than in mock-infected monolayers, we did not find a significant decrease in the number of Ca^2+^ spikes per FOV with CBX or ^10^Panx treatment (**Fig. 2C**). Furthermore, the dye loading scratch assay was also negative in jHIE monolayers (**Supplementary Fig. S1A**). Adding PGE2 or NOC-7 to jHIE-GCaMP6s cells also did not elicit a Ca^2+^ response (**Supplementary Fig. S1F-I**). These results demonstrate that RV-induced ICWs do not occur through gap junctions or *via* PGE2/NO in HIEs.

We used qPCR to determine the purinergic receptor expression in jHIE monolayers and found that P2×4, P2Y1, and P2Y2 had the highest expression, similar to that of MA104 cells (**Fig. 2G, 3D**). The relative mRNA expression levels of purinergic receptors did not substantially change with RV (Ito) infection (**Supplementary Fig. S2B**). To test the involvement of the different purinergic receptors in ICWs, we mock- or RV (Ito)-infected jHIE-GCaMP6s monolayers and treated with DMSO vehicle, apyrase, BPTU, AR-C 118925XX, Bx430, or 5BDBD, and we found that apyrase and BPTU treatment decreased the Ca^2+^ signaling and amplitude in RV-treated monolayers (**Fig. 3E, Supplementary Movie S7**). Quantification verified a significant decrease in the number of Ca^2+^ spikes/FOV and average Ca^2+^ spike magnitude with apyrase or BPTU treatment compared to the DMSO vehicle control in the RV-infected monolayers (**Fig. 3F**,**G**). Interestingly, BPTU and apyrase treatment also reduced the number of Ca^2+^ spikes in mock-inoculated monolayers (**Fig. 3F**), which indicates that jHIEs commonly utilize purinergic receptor-induced Ca^2+^ signaling under basal conditions and this is exploited during RV infection.

Since jHIEs are human-derived cultures, genetic variation may influence whether RV-infection can cause ICWs. We created an additional jHIE-GCaMP6s line derived from a different individual, and then mock- and RV-infected these jHIE-GCaMP6s monolayers and treated with a panel of P2Y1 receptor blockers (BPTU, MRS2179, MRS2279, and MRS2500). We found that all of the P2Y1 blockers significantly decreased the number of Ca^2+^ spike/FOV and average Ca^2+^ spike magnitude when compared to DMSO vehicle control for both the mock- or RV-infected monolayers (**Fig. 3H**,**I**). Together, these data support that RV-induced ICWs in intestinal epithelial cells occur by extracellular ADP signaling and the P2Y1 receptor, the same as in MA104 cells.

### CRISPR/Cas9 P2Y1 receptor knockout reduces intercellular calcium waves

Pharmacological studies of purinergic receptors have found that different receptors have distinct affinities for nucleotide binding, with the P2Y1 receptor having greater activation with ADP than with ATP (*37*). This suggests that ADP mediates the ICWs in RV infection, which is also consistent with studies that have found that RV-infected cells have reduced cytoplasmic concentrations of ATP (*38*). To test whether ADP is the signaling molecule of the ICWs, we infected MA104-GCaMP cells with RV (SA114F) and then treated with apyrase grade VI, which has a high ATPase/ADPase activity ratio, or apyrase grade VII, which has a low ATPase/ADPase activity ratio. Treatment with apyrase grade VII, but not grade VI, significantly decreased the number of Ca^2+^ spikes and the spike magnitude in RV-infected wells, supporting that ADP is the signaling molecule in RV-induced ICWs (**Fig. 4A**,**B**). To further test that the P2Y1 receptor is required for RV-induced Ca^2+^ waves, we genetically knocked out the P2Y1 receptor in MA104-GCaMP cells using lentivirus encoding CRISPR/Cas9 with small-guide RNAs targeted for the P2Y1 receptor gene (MA104-GCaMP-P2Y1ko). Genotyping analysis revealed several mutations in the sequence downstream of the guide RNA in 3 clonal populations of cells (**Supplementary Fig. S3A**,**B**). To test the P2Y1 knockout phenotype, we added ADP to parental and MA104-GCaMP-P2Y1ko cells and found a significantly decreased Ca^2+^ response in the knockout cells compared to the parental GECI line, consistent with a lack of P2Y1 receptor function (**Fig. 4C**).

**Figure 4.**
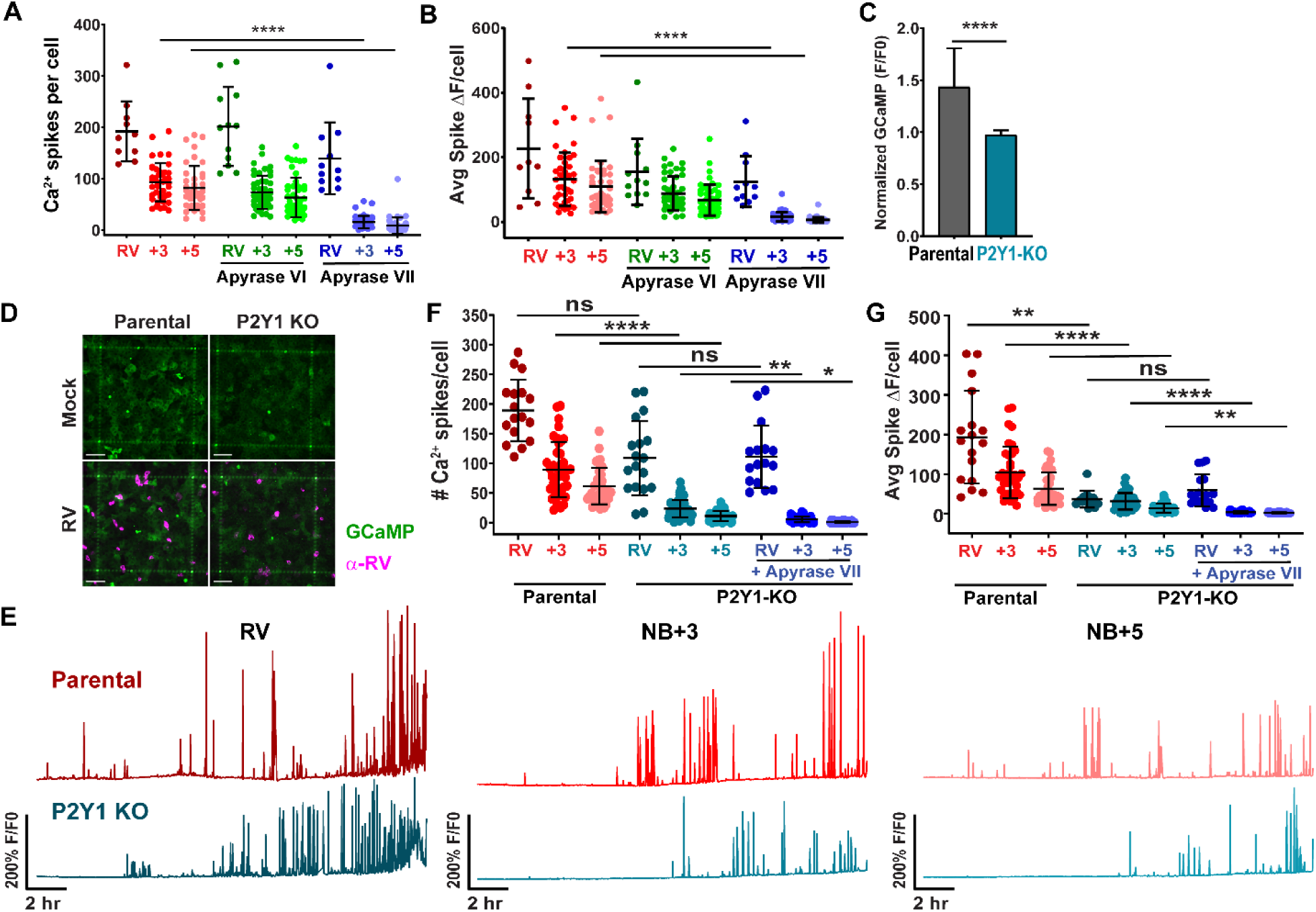
CRISPR/Cas9 knockout of P2Y1 receptor reduces intercellular calcium waves. **(A)** Ca^2+^ spikes and **(B)** average F/F_0_ increase of Ca^2+^ spikes in MA104-GCaMP cells infected with RV (SA114F) and treated with apyrase VI or VII. **(C)** F/F_0_ increase in parental or P2Y1 KO MA104-GCaMP cells treated with 1 nM ADP. Mann-Whitney test. **(D)** Immunofluorescence of parental or P2Y1ko cells mock- or RV-infected (pink). Scale bar = 100 µm **(E)** Ca^2+^ traces of parental or P2Y1ko cells infected with RV (SA11cl3-mRuby3). **(F)** Ca^2+^ spikes/cell in RV-infected or NB +3 or +5 cell and **(G)** the average F/F_0_ increase per spike. **(A-B, F-G)** Kruskal-Wallis with Dunn’s multiple comparisons test. (*p< 0.05, **p<0.01, ****p<0.0001)

Next, we sought to determine if knockout of the P2Y1 receptor would also prevent ICWs during RV infection. We mock- or RV (SA114F)-infected MA104-GCaMP (parental) or MA104-GCaMP-P2Y1ko cells. Fixing and immunostaining for RV antigen at 24 hpi showed positive staining for RV in parental and P2Y1 knockout cells confirming that the P2Y1 knockout did not prevent infection (**Fig. 4D**). In the P2Y1 knockout cells, there was a decrease in Ca^2+^ signaling, Ca^2+^ spikes, and average Ca^2+^ spike magnitude in the NB+3 and NB+5 cells compared with the parental, though not in the RV-infected cell (**Fig. 4E-G, Supplementary Movie S8**). Additional treatment of apyrase VII to the P2Y1 knockout cells only modestly decreased the Ca^2+^ spikes in neighboring cells (**Fig. 4G**).

Similarly, we tested the effect of genetic knockout of the P2Y1 receptor on ICWs in RV-infected jHIEs (jHIE-GCaMP6s-P2Y1ko). Genotyping analysis revealed two P2Y1 knockout enteroid populations comprised of an insertional mutation and a deletion in the sequence downstream of the guide RNA (**Supplementary Fig. S3C**,**D**). Phenotypic knockout was confirmed by a reduced Ca^2+^ response to ADP in jHIE-GCaMP6s-P2Y1ko enteroids compared to parental enteroids (**Fig. 5A**). For these studies we also created an additional jHIE-GCaMP6s enteroid line with CRISPR/Cas9 knockout of the P2Y2 receptor (jHIE-GCaMP6s-P2Y2ko). Genotyping analysis of the P2Y2ko enteroids detected two alleles with deletion mutations downstream of the guide RNA sequence (**Supplementary Fig. S4A**,**B**). To test the P2Y2 knockout phenotype, we added non-hydrolyzable ATP-γS to parental and P2Y2ko enteroids and found a significantly decreased Ca^2+^ response in the P2Y2ko jHIEs (**Fig. 5B**). Immunostaining the jHIE monolayers for RV antigen at 24 hpi showed positive staining for RV in parental, P2Y1ko, and P2Y2ko enteroids, demonstrating RV infection still occurred (**Fig. 5C**,**D**). Mock- or RV (Ito)- infection of the parental, jHIE-GCaMP6s-P2Y1ko, or jHIE-GCaMP6s-P2Y2ko resulted in significantly less Ca^2+^ signaling, Ca^2+^ spikes/FOV, and smaller average Ca^2+^ spike magnitude in the P2Y1 knockout jHIEs, but not in P2Y2 knockout jHIEs (**Fig. 5E-G, Supplementary Movie S9**). Interestingly, there also was significantly fewer Ca^2+^ spikes/FOV and average Ca^2+^ spike magnitude in the P2Y1 knockout enteroids compared to parental or P2Y2 knockout enteroids in mock-inoculation (**Fig. 5F**,**G**). Together, these data support that ADP and the P2Y1 receptor mediate the ICWs in RV infection and reduce basal ICW signaling.

**Figure 5.**
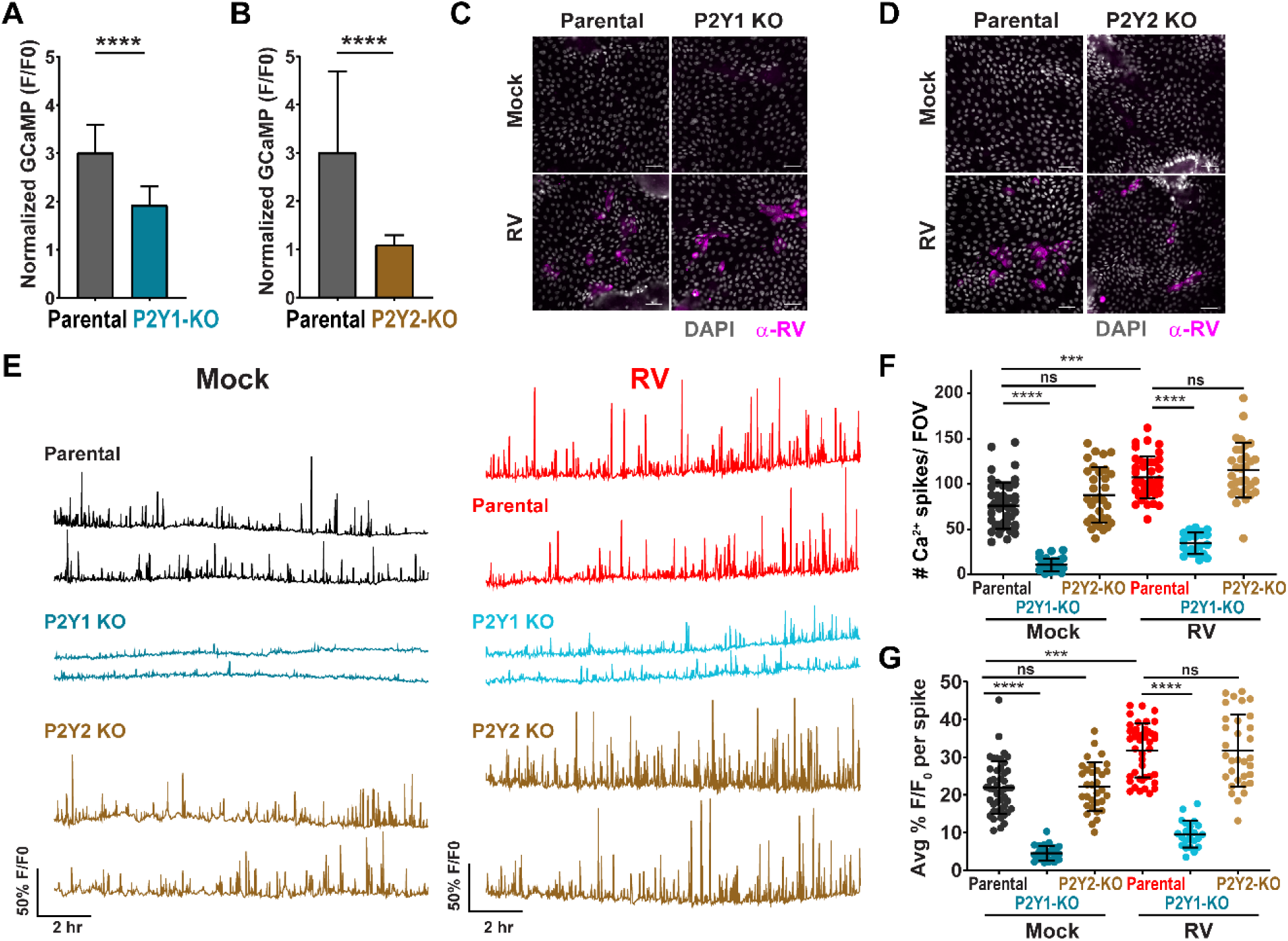
CRISPR/Cas9 knockout of P2Y1 receptor in jHIEs reduces intercellular calcium waves. **(A-B)** F/F_0_ increase in parental or **(A)** P2Y1ko (J2)HIE-GCaMP6s monolayers treated with 10 nM ADP and **(B)** P2Y2ko (J2)HIE-GCaMP6s monolayers treated with 10 µM ATP-γS. Mann-Whitney test. **(C-D)** Parental, P2Y1ko, or P2Y2ko jHIE-GCaMP6s monolayers mock- or RV(Ito)-infected, immunostained for RV antigen (pink). **(E)** Ca^2+^ traces/FOV of mock- or RV-infected **(F)** Ca^2+^ spikes/FOV and **(G)** average F/F_0_ increase. Mann-Whitney **(A-B)** or Kruskal-Wallis test with Dunn’s multiple corrections test **(F-G)**. (***p<0.001, ****p<0.0001) Scale bar = 50 µm.

### Purinergic signaling increases RV replication and expression of IL-1α, iNOS, and COX2

Previous studies have found that extracellular ATP reduces the replication of several viruses through activation of P2×7 receptors, which specifically bind ATP (*39, 40*). Less is known about the role of ADP in cellular signaling or its effects on viral infection and replication. We first tested the effect of purinergic blockers on RV replication. New infectious yield from RV (SA114F)-infected MA104 cells was decreased when treating cells with apyrase or BPTU after inoculation, and no significant decrease was observed when adding AR-C 118925XX, Bx430, 5BDBD, or the non-selective P2 receptor antagonists suramin and PPADS (**Fig. 6A**). Furthermore, RV infectious yield was also reduced from RV (SA11cl3-mRuby)-infected MA104-GCaMP-P2Y1ko cells compared to parental MA104-GCaMP cells (**Fig. 6B**). These results support that blocking ADP signaling and the P2Y1 receptor decreases RV replication.

**Figure 6.**
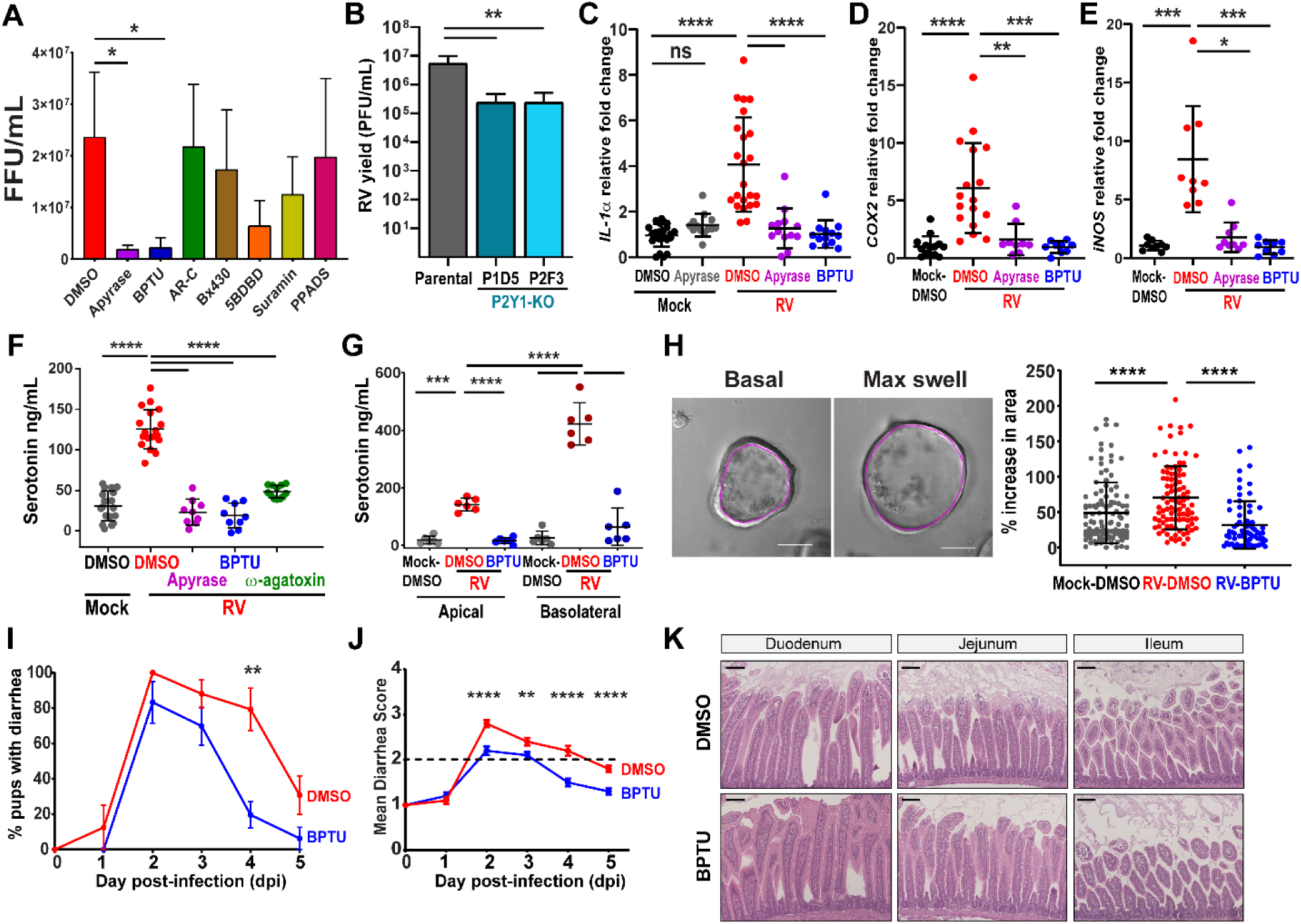
Purinergic signaling contributes to rotavirus disease processes. **(A)** Fluorescent focus yield of RV(SA114F)-infected MA104 cells. **(B)** Plaque yield of RV (SA114F)-infected MA104-GCaMP or MA104-GCaMP-P2Y1ko cells. **(C-E)** qPCR of mRNA transcripts in jHIE monolayers **(F-G)** Serotonin secretion from RV-infected jHIE **(F)** monolayers and **(G)** transwells. **(H)** Enteroid swelling assay using mock- or RV(Ito)-infected jHIEs treated with DMSO or 10 µM BPTU. **(I-K)** C57Bl/6J mouse pups with diarrhea infected with Rhesus RV and vehicle- or BPTU-treated and the **(J)** mean diarrhea score. Mean ± SEM. **(K)** Hematoxylin and eosin-stained intestinal sections, 5 dpi. Mann-Whitney **(H-J)**, Kruskal-Wallis with Dunn’s multiple comparisons **(C-E)** or one-way ANOVA with Bonferroni multiple comparisons test **(A-B, F-G).** (*p<0.05, **p<0.01, ***p<0.001, ****p<0.0001). Scale bar = 100 µm.

Extracellular ATP is also known to have pro-inflammatory functions and activate components of the innate immune system such as macrophages (*41, 42*). Previous studies have found that RV increases expression and release of the inflammatory cytokine interleukin-1α (IL-1α), which is highly expressed in epithelial cells (*43, 44*). Furthermore, strong cytosolic Ca^2+^ agonists can induce the secretion of IL-1α independent of inflammasome activation (*45*). To determine if RV-induced ICWs increase IL-1α, we mock- or RV (Ito)-infected jHIE monolayers and treated with apyrase or BPTU. RV infection significantly increased *IL-1α* expression over mock infection, and treatment with apyrase or BPTU inhibited this increase (**Fig. 6C**). This finding supports that RV-induced ICWs and ADP signaling increases *IL-1α* expression and thus mediates the inflammatory response from HIEs.

Since increased PGE2 and NO have been found during RV infection, we also investigated if RV-induced ICWs could stimulate their production (*17*–*19*). Previous studies have found that purinergic signaling stimulates cyclooxygenase-2 (*COX2*) expression and PGE2 release from Caco-2 and rat-derived IEC-6 cell lines and can also activate inducible nitric oxide synthase (*iNOS)* expression and nitric oxide production (*46, 47*). To test the role of ADP signaling, we mock- or RV (Ito)-infected jHIE monolayers and treated with apyrase or BPTU. RV infection significantly increased both *COX2* and *iNOS* expression over mock-infection, and treatment with apyrase or BPTU inhibited this increase (**Fig. 6D-E**). These findings suggest that both RV-induced ICWs and ADP signaling increase the expression of *COX2* and *iNOS* and thus potentially drive the increased PGE2 and NO observed in RV infections.

### Blocking purinergic signaling reduces serotonin and fluid secretion

RV infection causes a multifactorial secretory diarrhea, and two-thirds of the fluid secretion can be attributed to activation of the enteric nervous system (ENS) (*7*). Previous studies have found that RV infection can stimulate the release of serotonin from enterochromaffin cells to activate the ENS and blocking serotonin decreases diarrhea, making serotonin release a key mechanism for RV-induced diarrhea (*4, 48*). To determine if RV-induced ADP signaling could stimulate serotonin secretion, we mock- or RV (Ito)-infected jHIE monolayers and treated with DMSO vehicle, apyrase, or BPTU. After 24 hr, we found significantly increased serotonin in the supernatant of RV-infected jHIE monolayers compared to mock-infection, which was significantly inhibited by treatment with apyrase or BPTU (**Fig. 6F**). Furthermore, treatment of RV-infected monolayers with ω-agatoxin, a P-type voltage-gated Ca^2+^ channel blocker, also significantly inhibited serotonin levels, supporting that the measured serotonin was from cellular release rather than cell death (**Fig. 6F**). We found similar results when infecting jHIE transwells, a more polarized model of epithelium, with RV. Treatment with BPTU significantly decreased serotonin measured in both the apical and basolateral compartments compared with DMSO vehicle and overall a greater concentration of serotonin secreted basally (**Fig. 6G**). This data supports that ADP signaling stimulates serotonin secretion during RV infection.

RV infection also activates secretory pathways in epithelial cells through the stimulation of chloride secretion which drives paracellular movement of water. We tested whether RV-induced ADP signaling was important for RV-induced epithelial fluid secretion using the enteroid swelling assay, in which an increase in fluid secretion causes an increase in the internal lumen of three-dimensional (3D) enteroid cultures (*34, 49*). We mock- or RV-infected 3D jHIE-GCaMP6s enteroids, treated with DMSO vehicle or BPTU, and found that BPTU treatment significantly decreased the percentage of swelling in RV-infected enteroids (**Fig. 6H, Supplementary Movie S10**). This result supports that ADP signaling stimulates epithelial fluid secretion in RV infection.

Finally, we determined whether inhibition of the P2Y1 receptor would attenuate RV-induced diarrhea in a neonatal mouse model. BPTU consistently had the most potent inhibition of Ca^2+^ spikes and average Ca^2+^ spike magnitude in HIEs (**Fig. 3H**,**I**), and binds to an allosteric pocket on the external surface of the P2Y1 receptor (*50*). We tested a drug treatment model in which 6 to 8-day-old pups were orally gavaged with rhesus RV (RRV) and then gavaged with BPTU or vehicle control on days 1-4 post-infection. Stools were assessed each day and watery, mucus, yellow stools were indicators of diarrhea. Peak diarrhea occurred at 2 dpi in both DMSO- and BPTU-treated pups, but the percentage of pups with diarrhea decreased significantly on 4 dpi (**Fig. 6I**). Pups treated with BPTU also had significantly less severe diarrhea as measured by their mean diarrhea score (**Fig. 6J**). Furthermore, the intestinal cross-sections from the duodenum, jejunum, and ileum of RV-infected pups treated with DMSO or BPTU are relatively similar with no histopathological features at 5 dpi (**Fig. 6K**). These studies support that blocking the P2Y1 receptor decreased diarrhea severity and caused diarrhea to resolve more quickly than the untreated animals.

We next tested whether treatment with a water-soluble small molecule P2Y1 receptor blocker, MRS2500, would similarly attenuate RV diarrhea. While BPTU is an allosteric P2Y1 receptor antagonist, MRS2500 is a water soluble, nucleotide analog that binds to a pocket of the seven transmembrane domain bundle (*50*). In these studies, we found there were similar percentages of pups with diarrhea on dpi 1-5 (**Supplementary Fig. S5A**). However, pups treated with MRS2500 had significantly less severe diarrhea at the peak of diarrhea on dpi 3 and also on dpi4 compared to pups treated with saline (**Supplementary Fig. S5B**). These results further support that inhibition of the P2Y1 receptor decreases RV diarrhea severity.

## Discussion

The current concept of RV pathogenesis proposes that paracrine signaling from RV-infected cells dysregulates surrounding uninfected cells and contributes to life-threatening diarrhea and/or vomiting. Previous studies established that RV elevates cytosolic calcium during infection and increases paracrine signaling molecules such as eNSP4, PGE2, and NO, but functional signaling from RV-infected cells to uninfected cells had not been determined (*9, 18, 19*). Thus, the primary goal of this study was to investigate the role of intercellular signaling during RV infection and to identify the signaling mechanisms involved. Using GECI-expressing cell lines and HIEs, we used long-term live-cell fluorescence imaging to detect signaling between infected and uninfected cells. Our work describes the first direct observation of ICWs originating from RV-infected cells and has provided experimental proof for the widely-held paracrine signaling idea. However, the discovery of ADP signaling as both the paracrine signal and the most dominant Ca^2+^ signal observed in RV infection is paradigm-shifting. Extracellular purine signaling is a well-conserved communication system found in single-celled bacteria and protozoa as well as plants and higher order eukaryotes (*51*). RV is the first virus observed to activate ICWs and furthermore this signaling is mediated by ADP, rather than ATP, and the ADP-selective P2Y1 receptor. RV exploits ADP signaling in host cells to benefit its replication, but this signaling also leads to increased fluid and serotonin secretion that are hallmarks of RV disease. Furthermore, we found that treatment of neonatal mice with two mechanistically different small molecule P2Y1 receptor inhibitors was able to attenuate RV diarrhea, underlining that activation of purinergic signaling is biologically relevant to RV disease. Activating ICWs and ADP signaling may be a common strategy for enteric viruses to amplify dysregulation of host cells, and thus targeting the P2Y1 receptor may be an effective host-directed approach to treating viral diarrhea.

Additionally, ICWs and extracellular ADP signaling may be a broadly important form of communication in the gastrointestinal epithelium. Basal ICWs in jHIEs were reduced when treated with apyrase and P2Y1 blockers and P2Y1 receptor knockout enteroids had significantly reduced Ca^2+^ signaling. Since ATP-mediated ICWs are activated in wound healing responses (*52, 53*), tonic purinergic signaling may also be important for epithelial-to-epithelial cell communication and maintaining epithelial barrier integrity. Purinergic signaling is well known to regulate responses to injury and restitution by mediating the innate immune system (*42*). We speculate that basal, ADP-based ICWs may have a role in the balance of pro- and anti-inflammatory signaling from the epithelium. Furthermore, extracellular ATP signaling in the GI tract stimulates epithelial ion transport (*54*), suggesting a role for basal ICWs to stimulate fluid secretion and hydrate the mucus layer. Finally, ADP-driven ICWs may mediate communication between the epithelium to the enteric nervous system directly or signal to the lamina propria and endothelium as part of tissue-level homeostasis. As the epithelium is the primary interface between the environment and the host, understanding how it coordinates and communicates is critical for developing therapeutics when it becomes dysregulated in disease states.

Together, these studies identify ADP as the signaling compound that elicits a paracrine signal involved in RV diarrhea and/or vomiting responses. Our studies demonstrated that reduction of ADP signaling and ICWs decreases many known aspects of RV disease including serotonin release, epithelial fluid secretion, and diarrhea in a neonatal mouse model. Since ICWs and ADP signaling are involved in RV disease processes, targeting the P2Y1 receptor may be an effective approach to treating viral diarrhea. This is the first known example that viruses can exploit purinergic signaling and ICWs, and this may be a common strategy for enteric viruses to amplify dysregulation of pathophysiological signaling important for diarrhea.

## Acknowledgements

This work was supported in part by NIH grants K01DK093657, R03DK110270, R01DK115507 (PI: J. M. Hyser), and R01AI080656 and U19AI116497 (PI: M. K. Estes). Trainee support for A.C.G. was provided by NIH grants F30DK112563 (PI: A. Chang-Graham) and the BCM Medical Scientist Training Program and support for both A.C.G. and A.C.S. was provided by the Integrative Molecular and Biomedical Sciences Graduate Program (T32GM008231, PI: D. Nelson). This project was supported in part by PHS grant P30DK056338, which supports the Texas Medical Center Digestive Diseases Center (TMC-DDC) Gastrointestinal Experimental Model Systems (GEMS) Core and the Cellular and Molecular Morphology Core. We would like to thank Xi-Lei (Shelly) Zeng and Xiaomin Yu for their help with enteroid cultures and media. FACS sorting of cell lines utilized the BCM Cytometry and Cell Sorting Core with funding from the CPRIT Core Facility Support Award (CPRIT-RP180672), the NIH (CA125123 and RR024574), and the expert assistance of Joel M. Sederstrom. Additional thanks to the BCM Center for Comparative Medicine and technicians for their care of the animals.

## Author Contributions

JH, ACG, JLP, MAE, HD, and MKE designed the experiments and discussed the data. JH, ACG, JLP, FS, AS, JN, and JK conducted the calcium imaging experiments and analyzed the data with JH and ACG. JLP conducted the fluorescent focus assays, plaque assays, and genotyping and analyzed data with JH. JLP and ACG created the CRISPR/Cas9 knockout lines. MAE and KE conducted and analyzed qPCR experiments with ACG. HD performed and analyzed the serotonin ELISAs. ACG conducted and analyzed the enteroid swelling assays. JH and ACG performed the mouse experiments and MAE performed immunohistochemistry. NP and MKE provided key reagents including the anti-NSP4 antibodies. JH and ACG wrote the manuscript, and all authors contributed to revisions of the paper.

## Data and materials availability

All data is available in the manuscript or the supplementary materials. The data that support the findings of this study are available from the corresponding author upon reasonable request. Correspondence and requests for materials should be addressed to Joseph Hyser

## References

1. C. Troeger, I. A. Khalil, P. C. Rao, S. Cao, B. F. Blacker, T. Ahmed, G. Armah, J. E. Bines, T. G. Brewer, D. V. Colombara, G. Kang, B. D. Kirkpatrick, C. D. Kirkwood, J. M. Mwenda, U. D. Parashar, W. A. Petri, M. S. Riddle, A. D. Steele, R. L. Thompson, J. L. Walson, J. W. Sanders, A. H. Mokdad, C. J. L. Murray, S. I. Hay, R. C. Reiner, Rotavirus Vaccination and the Global Burden of Rotavirus Diarrhea among Children Younger Than 5 Years. JAMA Pediatr. 172, 958–965 (2018).

2. G. P. Davidson, I. Goller, R. F. Bishop, R. R. Townley, I. H. Holmes, B. J. Ruck, Immunofluorescence in duodenal mucosa of children with acute enteritis due to a new virus. J. Clin. Pathol. 28, 263–266 (1975).

3. W. G. Starkey, J. Collins, T. S. Wallis, G. J. Clarke, A. J. Spencer, S. J. Haddon, M. P. Osborne, D. C. Candy, J. Stephen, Kinetics, tissue specificity and pathological changes in murine rotavirus infection of mice. J. Gen. Virol. 67, 2625–2634 (1986).

4. M. Hagbom, C. Istrate, D. Engblom, T. Karlsson, J. Rodriguez-Diaz, J. Buesa, J. a. Taylor, V. M. Loitto, K. E. Magnusson, H. Ahlman, O. Lundgren, L. Svensson, Rotavirus stimulates release of serotonin (5-HT) from human enterochromaffin cells and activates brain structures involved in nausea and vomiting. PLoS Pathog. 7, 1–10 (2011).

5. J. A. Little, Lynn M. Shadduck, Pathogenesis of rotavirus infection. Infect. Immun. 38, 755–763 (1982).

6. A. P. Morris, M. K. Estes, VIII. Pathological consequences of rotavirus infection and its enterotoxin. Am. J. Physiol. Liver Physiol. 281, G303–G310 (2001).

7. O. Lundgren, a T. Peregrin, K. Persson, S. Kordasti, I. Uhnoo, L. Svensson, Role of the enteric nervous system in the fluid and electrolyte secretion of rotavirus diarrhea. Science. 287, 491–495 (2000).

8. J. M. Hyser, M. R. Collinson-Pautz, B. Utama, M. K. Estes, Rotavirus disrupts calcium homeostasis by NSP4 viroporin activity. MBio. 1, 1–12 (2010).

9. J. M. Ball, P. Tian, C. Q. Zeng, a P. Morris, M. K. Estes, Age-dependent diarrhea induced by a rotaviral nonstructural glycoprotein. Science. 272, 101–104 (1996).

10. P. Tian, M. K. Estes, Y. Hu, J. M. Ball, C. Q. Zeng, W. P. Schilling, The rotavirus nonstructural glycoprotein NSP4 mobilizes Ca2+ from the endoplasmic reticulum. J. Virol. 69, 5763–5772 (1995).

11. K. Kunzelmann, R. Schreiber, A. Kmit, W. Jantarajit, J. R. Martins, D. Faria, P. Kongsuphol, J. Ousingsawat, Y. Tian, Expression and function of epithelial anoctamins. Exp. Physiol. 285, 184–192 (2011).

12. S. E. Crawford, J. M. Hyser, B. Utama, M. K. Estes, Autophagy hijacked through viroporin-activated calcium/calmodulin-dependent kinase kinase-β signaling is required for rotavirus replication. Proc. Natl. Acad. Sci. U. S. A. 109, E3405–13 (2012).

13. J. L. Zambrano, O. Sorondo, A. Alcala, E. Vizzi, Y. Diaz, M. C. Ruiz, F. Michelangeli, F. Liprandi, J. E. Ludert, Rotavirus Infection of Cells in Culture Induces Activation of RhoA and Changes in the Actin and Tubulin Cytoskeleton. PLoS One. 7, e47612 (2012).

14. I. Beau, J. Cotte-Laffitte, R. Amsellem, A. L. Servin, A protein kinase A-dependent mechanism by which rotavirus affects the distribution and mRNA level of the functional tight junction-associated protein, occludin, in human differentiated intestinal Caco-2 cells. J. Virol. 81, 8579–8586 (2007).

15. M. Hagbom, S. Sharma, O. Lundgren, L. Svensson, Towards a human rotavirus disease model. Curr. Opin. Virol. 2, 408–418 (2012).

16. K. Hodges, R. Gill, Infectious diarrhea: Cellular and molecular mechanisms. Gut Microbes. 1, 4–21 (2010).

17. T. V. Sowmyanarayanan, S. K. Natarajan, A. Ramachandran, R. Sarkar, P. D. Moses, A. Simon, I. Agarwal, S. Christopher, G. Kang, Nitric oxide production in acute gastroenteritis in Indian children. Trans. R. Soc. Trop. Med. Hyg. 103, 849–51 (2009).

18. Y. Yamashiro, T. Shimizu, S. Oguchi, M. Sato, Prostaglandins in the plasma and stool of children with rotavirus gastroenteritis. J. Pediatr. Gastroenterol. Nutr. 9, 322–7 (1989).

19. J. Rodriguez-Diaz, M. Banasaz, C. Istrate, J. Buesa, O. Lundgren, F. Espinoza, T. Sundqvist, M. Rottenberg, L. Svensson, Role of Nitric Oxide During Rotavirus Infection. J. Med. Virol. 78, 979–985 (2006).

20. A. P. Morris, J. K. Scott, J. M. Ball, C. Q.-Y. Zeng, W. K. O’Neal, M. K. Estes, NSP4 elicits age-dependent diarrhea and Ca2+-mediated I-influx into intestinal crypts of CF mice. Am. J. Physiol. Gastrointest. Liver Physiol. 277, 431–444 (1999).

21. J. Ousingsawat, M. Mirza, Y. Tian, E. Roussa, R. Schreiber, D. I. Cook, K. Kunzelmann, Rotavirus toxin NSP4 induces diarrhea by activation of TMEM16A and inhibition of Na+ absorption. Pflugers Arch. Eur. J. Physiol. 461, 579–589 (2011).

22. N. Seo, C. Q. Zeng, J. M. Hyser, B. Utama, S. E. Crawford, K. J. Kim, M. Ho, M. K. Estes, Integrins are receptors for the rotavirus enterotoxin. Proc. Natl. Acad. Sci. 105, 8811–8818 (2008).

23. S. Fujii, K. Suzuki, A. Kawamoto, F. Ishibashi, T. Nakata, T. Murano, G. Ito, H. Shimizu, T. Mizutani, S. Oshima, K. Tsuchiya, T. Nakamura, A. Araki, K. Ohtsuka, R. Okamoto, M. Watanabe, PGE 2 is a direct and robust mediator of anion/fluid secretion by human intestinal epithelial cells. Sci. Rep. 6, 1–15 (2016).

24. F. H. Mourad, J. L. Turvill, M. J. G. Farthing, Role of nitric oxide in intestinal water and electrolyte transport. Gut. 44, 143–147 (1999).

25. F. Michelangeli, M. C. Ruiz, J. R. del Castillo, J. E. Ludert, F. Liprandi, Effect of rotavirus infection on intracellular calcium homeostasis in cultured cells. Virology. 181, 520–527 (1991).

26. J. M. Hyser, M. K. Estes, Annu. Rev. Virol., in press, doi:10.1146/annurev-virology-100114-054846.

27. S. E. Crawford, S. Ramani, J. E. Tate, U. D. Parashar, L. Svensson, M. Hagbom, M. A. Franco, H. B. Greenberg, M. O’Ryan, G. Kang, U. Desselberger, M. K. Estes, Rotavirus infection. Nat. Rev. Dis. Prim. 3 (2017), doi:10.1038/nrdp.2017.83.

28. A. L. Chang-Graham, J. L. Perry, A. C. Strtak, N. K. Ramachandran, J. M. Criglar, A. A. Philip, J. T. Patton, M. K. Estes, J. M. Hyser, Rotavirus Calcium Dysregulation Manifests as Dynamic Calcium Signaling in the Cytoplasm and Endoplasmic Reticulum. Sci. Rep. 9 (2019), doi:10.1101/627877.

29. M. J. Sanderson, A. C. Charles, E. R. Dirksen, Mechanical stimulation and intercellular communication increases intracellular Ca2+ in epithelial cells. Cell Regul. 1, 585–96 (1990).

30. A. H. Cornell-Bell, S. M. Finkbeiner, M. S. Cooper, S. J. Smith, Glutamate induces calcium waves in cultured astrocytes: long-range glial signaling. Science. 247, 470–3 (1990).

31. L. Leybaert, M. J. Sanderson, Intercellular Ca(2+) waves: mechanisms and function. Physiol. Rev. 92, 1359–1392 (2012).

32. K. R. Emslie, J. M. Miller, M. B. Slade, P. R. Dormitzer, H. B. Greenberg, K. L. Williams, Expression of the rotavirus SA11 protein VP7 in the simple eukaryote Dictyostelium discoideum. J. Virol. 69, 1747–1754 (1995).

33. T. Sato, D. E. Stange, M. Ferrante, R. G. J. Vries, J. H. Van Es, S. Van Den Brink, W. J. Van Houdt, A. Pronk, J. Van Gorp, P. D. Siersema, H. Clevers, Long-term expansion of epithelial organoids from human colon, adenoma, adenocarcinoma, and Barrett’s epithelium. Gastroenterology. 141, 1762–1772 (2011).

34. K. Saxena, S. E. Blutt, K. Ettayebi, X. Zeng, J. R. Broughman, S. E. Crawford, U. C. Karandikar, N. P. Sastri, M. E. Conner, A. R. Opekun, D. Y. Graham, W. Qureshi, V. Sherman, J. Foulke-Abel, J. In, O. Kovbasnjuk, N. C. Zachos, M. Donowitz, M. K. Estes, Human Intestinal Enteroids: a New Model To Study Human Rotavirus Infection, Host Restriction, and Pathophysiology. J. Virol. 90, 43–56 (2016).

35. J. Foulke-Abel, J. In, O. Kovbasnjuk, N. C. Zachos, K. Ettayebi, S. E. Blutt, J. M. Hyser, X.-L. Zeng, S. E. Crawford, J. R. Broughman, M. K. Estes, M. Donowitz, Human enteroids as an ex-vivo model of host-pathogen interactions in the gastrointestinal tract. Exp. Biol. Med. 239, 1124–1134 (2014).

36. N. C. Zachos, O. Kovbasnjuk, J. Foulke-Abel, J. In, S. E. Blutt, H. R. De Jonge, M. K. Estes, M. Donowitz, Human enteroids/colonoids and intestinal organoids functionally recapitulate normal intestinal physiology and pathophysiology. J. Biol. Chem. 291, 3759–3766 (2016).

37. C. Léon, B. Hechler, C. Vial, C. Leray, J. P. Cazenave, C. Gachet, The P2Y1 receptor is an ADP receptor antagonized by ATP and expressed in platelets and megakaryoblastic cells. FEBS Lett. 403, 26–30 (1997).

38. K. G. Dickman, S. J. Hempson, J. Anderson, S. Lippe, L. Zhao, R. Burakoff, R. D. Shaw, Rotavirus alters paracellular permeability and energy metabolism in Caco-2 cells. Am. J. Physiol. Gastrointest. Liver Physiol. 279, G757–G766 (2000).

39. C. Zhang, H. He, L. Wang, N. Zhang, H. Huang, Q. Xiong, Y. Yan, N. Wu, H. Ren, H. Han, M. Liu, M. Qian, B. Du, Virus-Triggered ATP Release Limits Viral Replication through Facilitating IFN-β Production in a P2×7-Dependent Manner. J. Immunol. 199, 1372–1381 (2017).

40. D. Ferrari, M. Idzko, T. Müller, R. Manservigi, P. Marconi, Purinergic Signaling: A New Pharmacological Target Against Viruses? Trends Pharmacol. Sci. 39, 926–936 (2018).

41. S. Zumerle, B. Calì, F. Munari, R. Angioni, F. Di Virgilio, B. Molon, A. Viola, Intercellular Calcium Signaling Induced by ATP Potentiates Macrophage Phagocytosis. Cell Rep. 27, 1-10.e4 (2019).

42. C. Cekic, J. Linden, Purinergic regulation of the immune system. Nat. Rev. Immunol. 16, 177–192 (2016).

43. P. P. Hernández, T. Mahlakõiv, I. Yang, V. Schwierzeck, N. Nguyen, F. Guendel, K. Gronke, B. Ryffel, C. Hölscher, L. Dumoutier, J. C. Renauld, S. Suerbaum, P. Staeheli, A. Diefenbach, Interferon-γ and interleukin 22 act synergistically for the induction of interferon-stimulated genes and control of rotavirus infection. Nat. Immunol. 16, 698–707 (2015).

44. N. C. Di Paolo, D. M. Shayakhmetov, Interleukin 1α and the inflammatory process. Nat. Immunol. 17, 906–913 (2016).

45. O. Groß, A. S. Yazdi, C. J. Thomas, M. Masin, L. X. Heinz, G. Guarda, M. Quadroni, S. K. Drexler, J. Tschopp, Inflammasome Activators Induce Interleukin-1α Secretion via Distinct Pathways with Differential Requirement for the Protease Function of Caspase-1. Immunity. 36, 388–400 (2012).

46. Y. Ohtani, M. Minami, M. Satoh, Expression of inducible nitric oxide synthase mRNA and production of nitric oxide are induced by adenosine triphosphate in cultured rat microglia. Neurosci. Lett. 293, 72–74 (2000).

47. E. Degagné, D. M. Grbic, A.-A. Dupuis, E. G. Lavoie, C. Langlois, N. Jain, G. A. Weisman, J. Sévigny, F.-P. Gendron, P2Y 2 Receptor Transcription Is Increased by NF-κB and Stimulates Cyclooxygenase-2 Expression and PGE 2 Released by Intestinal Epithelial Cells. J. Immunol. 183, 4521–4529 (2009).

48. S. Bialowas, M. Hagbom, J. Nordgren, T. Karlsson, S. Sharma, K.-E. Magnusson, L. Svensson, Rotavirus and Serotonin Cross-Talk in Diarrhoea. PLoS One. 11, e0159660 (2016).

49. J. F. Dekkers, C. L. Wiegerinck, H. R. de Jonge, I. Bronsveld, H. M. Janssens, K. M. Winter-de Groot, A. M. Brandsma, N. W. M. de Jong, M. J. C. Bijvelds, B. J. Scholte, E. E. S. Nieuwenhuis, S. van den Brink, H. Clevers, C. K. van der Ent, S. Middendorp, J. M. Beekman, A functional CFTR assay using primary cystic fibrosis intestinal organoids. Nat. Med. 19, 939–945 (2013).

50. D. Zhang, Z. G. Gao, K. Zhang, E. Kiselev, S. Crane, J. Wang, S. Paoletta, C. Yi, L. Ma, W. Zhang, G. W. Han, H. Liu, V. Cherezov, V. Katritch, H. Jiang, R. C. Stevens, K. A. Jacobson, Q. Zhao, B. Wu, Two disparate ligand-binding sites in the human P2Y1 receptor. Nature. 520, 317–321 (2015).

51. A. Verkhratsky, G. Burnstock, Biology of purinergic signalling: Its ancient evolutionary roots, its omnipresence and its multiple functional significance. BioEssays. 36, 697–705 (2014).

52. H. Takada, K. Furuya, M. Sokabe, Mechanosensitive ATP release from hemichannels and Ca 2 + influx through TRPC6 accelerate wound closure in keratinocytes. J. Cell Sci. 127, 4159–4171 (2014).

53. Y. Sung, Z. Sung, C. Ho, M. Lin, J. Wang, S. Yang, Y. Chen, C. Lin, Intercellular calcium waves mediate preferential cell growth toward the wound edge in polarized hepatic cells. Exp. Cell Res. 287, 209–218 (2003).

54. I. Novak, Purinergic signalling in epithelial ion transport: regulation of secretion and absorption. Acta Physiol. 202, 501–522 (2011).

