## Supplemental Methods and Figures for "Rotavirus Induces Intercellular Calcium Waves through ADP Signaling"

**This PDF file includes:**

Materials and Methods
Figs. S1 to S5
Tables S1 to S2
Captions for Movies S1 to S10

**Other Supplementary Materials for this manuscript include the following:**

Movies S1 to S10

### Materials and Methods

**Cells and rotaviruses.** MA104 cells (African green monkey kidney cells) and HEK293T cells (ATCC CRL-3216) were cultured in high glucose DMEM supplemented with 10% fetal bovine serum (FBS) and Antibiotic/Antimycotic (Invitrogen) at 37°C in 5% $CO_2$. The rotavirus human strain Ito, used for all human intestinal enteroid infections, was prepared as previously described (*34*). Rotavirus strains SA114F and RRV were produced from in-house stocks. Recombinant and SA11 clone 3-mRuby3 was produced previously (*28*). All viruses were propagated in MA104 cells in serum-free DMEM supplemented with 1 μg/mL Worthington's Trypsin, and after harvest stocks were subjected to three freeze/thaw cycles and activated with 10 μg/mL Worthington's Trypsin for 30 min at 37 °C prior to use.

**Rotavirus diarrhea mouse model.** C57Bl/6J dams with natural litters (purchased from Center for Comparative Medicine, Baylor College of Medicine) were used and housed in standard cages with food and water provided *ad libitum*. Mouse pups, both male and female at natural ratios, were housed with their dam throughout the experiment. Animal experiments were approved by the Baylor College of Medicine Institutional Animal Care and Use Committee (IACUC), approved protocol AN-6903. All experiments were performed in accordance with the recommendations in the Guide for the Care and Use of Laboratory Animals of the National Institutes of Health.

Six- or seven-day-old mouse pups were orally gavaged with 1.5 x $10^7$ PFU/mL of Rhesus rotavirus, a dosage sufficient to cause diarrhea in 100% of pups. The inoculum was mixed with sterile-filtered green food dye (3% by volume) for visualization of the gavage. For BPTU experiments, pups were orally gavaged with BPTU (4 mg/kg) (4 cages, n = 30 pups) or an equivalent volume of DMSO vehicle control (4 cages, n = 26 pups) suspended in pharmaceutical grade corn oil on days 1-4 post-infection. For MRS2500 experiments, pups were orally gavaged with MRS2500 (4 mg/kg) (9 cages, n = 60 pups) suspended in pharmaceutical grade normal saline or an equivalent volume of saline control (8 cages, n = 46 pups) on days 1-4 post-infection.

Each day, mouse pups were gently palpated on the abdomen to express and visualize feces. Stools were assessed on a scale between 1-4 for volume, color, and consistency as described previously (*9*), an average score of ≥2 was considered diarrhea. For scoring consistency, select experiments were scored by an observer blinded to experimental condition. Percentage of pups with diarrhea was calculated as the number of stools with a diarrhea score ≥ 2 divided by the number of stools collected per cage per day. Pups from which a stool sample could not be collected on that day were not counted for percent diarrhea or mean diarrhea severity calculations.

**Chemicals.** BPTU, AR-C 118925XX, Bx430, 5-BDBD, MRS2179, MRS2279, MRS2500, 10-Panx, suramin, TAT-Gap19, PPADS (Pyridoxalphosphate-6-azophenyl-2',4'-disulfonic acid), NOC-7, prostaglandin E2, and ω-agatoxin were purchased from Tocris Bioscience. Apyrase grade VI (catalog no. A6410) and apyrase grade VII (catalog no. A6535) were purchased from Sigma-Aldrich. 18β-glycyrrhetinic acid and carbenoxolone were purchased from Santa Cruz Biotechnology.

**Antibodies.** To detect RV by IF, we used rabbit anti-RV strain Alabama (*12*) (1:80,000) and the secondary antibody donkey anti-rabbit AlexaFluor 568 (Invitrogen) (1:2000). For eNSP4-blocking assays, we used purified anti-NSP4 rabbit antisera (120-147)(*22*), purified anti-NSP4 monoclonal antibody 622 (data in preparation), and purified M60, a non-neutralizing anti-VP7 monoclonal antibody, was used as a control (*32*). Equal concentrations of antibody, estimated using protein OD280, were added to RV-infected wells after inoculation.

**Calcium indicator lentiviruses, cell lines, and enteroids**. GCaMP5G (Addgene plasmid #31788), GCaMP6s (Addgene plasmid #40753), and G-CEPIA1er (Addgene plasmid #105012) were cloned into pLVX-Puro. RGECO1.2 (Addgene plasmid #45494), R-CEPIA1er (Addgene plasmid #58216), and GCaMP6s were cloned into pLVX-IRES-Hygro. Lentivirus vectors for the GECI constructs were packaged in HEK293T cells as previously described(*55*) or produced commercially (Cyagen Biosciences, Inc.). MA104-GCaMP5G, MA104-GCaMP5G/RCEPIAer, MA104-RGECO1/GCEPIAer, and MA104-GCaMP6s-shRNA cell lines and the jHIE-GCaMP6s enteroids (enteroid line J3) were generated as previously described (*28*, *55*). Both MA104-GCaMP5G and MA104-GCaMP6s cells were used in these studies; we observed no difference in calcium responses with these two indicators and thus refer to both as MA104-GCaMP. Jejunum (J2) human intestinal enteroid cultures expressing GCaMP6s (in pLVX-IRES-Hygro) were created using lentivirus transduction as described previously and grown in high Wnt3a CMGF+ with 50 µg/mL hygromycin B for selection (*56*).

**CRISPR/Cas9 KO cell lines and enteroids.** Lentivirus constructs in pLentiCRISPRv2 vectors were purchased from GenScript Biotech Corp (USA) and were packaged in HEK293T cells as previously described (*55*). The CRISPR/Cas9-expression vectors encode a puromycin-resistance gene for drug selection. The following small guide RNAs were used: P2Y1-sg1 CTACAGCATGTGCACGACCG and P2Y2-sg1 TTTGTCACCACCAGCGCGCG. MA104-GCaMP6s cells (pLVX-IRES-Hygro) were transduced with the P2Y1 CRISPR/Cas9-expressing construct and at 72 hrs post-transduction the cells were passaged in the presence of 50 µg/mL hygromycin B and 10 µg/mL puromycin to select for co-expression of GCaMP6s and CRISPR/Cas9. The J2-derived jHIE-GCaMP6s(hygro) enteroid cultures described above were transduced with the CRISPR/Cas9-expressing constructs to make two separate enteroid lines and grown with 50 μg/mL hygromycin B and 1 μg/mL puromycin to select for co-expression of GCaMP6s and CRISPR/Cas9.

Genotyping was performed on whole genomic DNA prepared using DirectPCR (Viagen) with Proteinase K (Qiagen) according to manufacturer's instructions. Genomic DNA was a template to PCR amplify a region surrounding the P2Y1 or P2Y2 sgRNA-target sites using primers listed in **Table S1**. PCR products were cloned into the pMiniT2.0 vector using the NEB PCR Cloning Kit (New England Biolabs). Five clones for P2Y1ko HIEs and five clones for P2Y2ko HIEs were picked and sequenced to determine the repertoire of alleles present in the HIE cultures. Sanger sequencing was performed by GENEWIZ (USA).

Phenotypic knockout was tested through measurement of relative GFP fluorescence increase after addition of 1 nM ADP for MA104-GCaMP6s-P2Y1ko cells, 10 nM ADP for (J2)HIE-GCaMP6s-P2Y1ko enteroids and 10 μM ATP-γS for (J2)HIE-GCaMP6s-P2Y2ko enteroids compared to parental (J2)HIE-GCaMP6s-P2Y2ko enteroids. Experiments were performed at least 3 times with $n \geq 25$ cells per experiment analyzed.

**Establishment of HIE cultures.** The human intestinal enteroid (HIE) cultures used in this study were generated previously and deposited into the Texas Medical Center Digestive Diseases Center (TMC-DDC) Gastrointestinal Experimental Model Systems (GEMS) Core (*33*, *34*). We used jejunal HIEs cultures J2 and J3, both from adult females, obtained from the GEMS core. Complete media with and without growth factors (CMGF+ and CMGF-, respectively) (*33*, *34*), differentiation media (*56*), high Wnt3a CMGF+ (hW-CMGF+) (*56*), and a Fluorobrite-DMEM based differentiation media (FB-Diff) were prepared as previously described (*28*). HIEs were grown in phenol red-free, growth factor-reduced Matrigel (Corning). HIE monolayers were prepared from three-dimensional cultures and seeded into optical-bottom 10-well Cellview chamber slides coated with dilute collagen IV (Sigma) as described previously (*57*, *58*). After 24 hr in CMGF+ and 10 µM Y-27632 Rock inhibitor, differentiation medium was used and changed every day for 4-5 days.

**Microscopy and image analysis.** For calcium imaging, MA104 cells and HIEs were imaged with a widefield epifluorescence Nikon TiE inverted microscope using a SPECTRAX LED light source (Lumencor) and either a 20x Plan Fluor (NA 0.45) phase contrast or a 20X Plan Apo (NA 0.75) differential interference contrast (DIC) objective. Fluorescence and transmitted light images were recorded using an ORCA-Flash 4.0 sCMOS camera (Hamamatsu), and Nikon Elements Advanced Research v4.5 software was used for multipoint position selection, data acquisition, and image analysis. Images were read-noise subtracted using an average of 10 no-light acquisitions of the camera. Single cells were selected as Regions of Interest (ROI) and fluorescence intensity measured for the experiment. The fluorescence intensity of the whole field-of-view was measured for HIE monolayers. Fluorescence intensity values were exported to Microsoft Excel and normalized to the baseline fluorescence. The number and magnitude of $Ca^{2+}$ spikes were calculated as described previously (*28*). $Ca^{2+}$ signals with a $\Delta F$ magnitude of > 5% were counted as $Ca^{2+}$ spikes. Average relative fluorescence increase in $Ca^{2+}$ spike magnitude was calculated from the highest 50 $Ca^{2+}$ spikes per cell or per field of view for HIE monolayers.

**Calcium imaging.**

*MA104-GECI cells.* Fluorobrite DMEM supplemented with 15mM HEPES, 1X sodium pyruvate, 1X Glutamax, and 1X non-essential amino acids (Invitrogen) was used for fluorescence $Ca^{2+}$ imaging (FB-Plus). Confluent monolayers of MA104-GCaMP cells in gridded 8-well chamber slides (ibidi) were mock- or RV-infected in FBS-free media for 1 hr at the indicated multiplicity of infection (MOI). The inoculum was removed and replaced with FB-Plus, and for experiments where noted, with DMSO vehicle or drugs at indicated concentrations. For experiments with gap junction blockers, either 30 µM 18β-glycyrrhetinic acid (18β-gly) or 50 µM TAT-Gap19 (Gap19) was added. For experiments with apyrase, either 10 U/mL of apyrase VI, 10 U/mL of apyrase VII, or a mixture of 5 U/mL of apyrase VI and 5 U/mL of apyrase VII was added. For experiments with purinergic receptor blockers, 10 μM BPTU, 10 μM AR-C 118925XX, 10 μM Bx430, or 10 μM 5-BDBD was used or otherwise noted. The slide was mounted into an Okolab microscope stage-top incubation chamber equilibrated to 37°C with a humidified 5% $CO_2$ atmosphere. In each experiment, 3-5 positions per well were selected and imaged every 1 minute for ~3-22 hours post infection. Each imaging experiment was performed at least three times independently, and data shown are representative of 1-3 experiments combined.

*GECI HIEs.* For RV infection in monolayers, jHIE-GCaMP6s monolayers were washed once with CMGF- and treated with an inoculum of 50 μL CMGF- plus 30 μL MA104 cell lysate or RV (Ito) and incubated at 37°C and 5% $CO_2$ for 2 hr. Then inoculum was removed, and monolayers were washed once with FB-Diff before adding FB-Diff with DMSO or drugs at indicated concentrations. For experiments with apyrase, a mixture of 50 U/mL of apyrase VI and 50 U/mL of apyrase VII was added. For experiments with purinergic receptor blockers, 10 μM BPTU, 10 μM AR-C 118925XX, 10 μM Bx430, 10 μM 5-BDBD, 10 μM MRS2179, 10 μM MRS2279, or 10 μM MRS2500 was used. Monolayers were transferred to the stage-top incubator for imaging with 4 fields of view chosen per well and imaged every minute for ~7-22 hours post-infection. Each imaging experiment was performed at least three times independently, and data shown are representative of all replicates combined.

**Enteroid swelling assay.** Jejunum HIEs were split and grown in hW-CMGF+ for 2 days followed by differentiation medium for 1 day. jHIEs were gently washed using ice cold 1XPBS and suspended in inoculum of 50 μL MA104 cell lysate or RV (Ito) diluted with 150 μL CMGF- and incubated for 1 hr. Then HIEs were washed, resuspended in 25% Matrigel diluted in FB-Diff (with DMSO vehicle or BPTU) and pipetted onto 8-well chamber slides (Matek) pre-coated with

Matrigel. Imaging positions were chosen so that between 30-50 enteroids were selected per experimental condition. Enteroids were imaged using the stage-top incubator with transmitted light and GFP fluorescence every 2-3 minutes for ~18 hrs. Swelling was determined by measuring the cross-sectional area of the internal lumen at the beginning of the run compared to the time point of maximum swelling and calculating the percent increase. Area measurements were determined using the Nikon Elements AR v4.5 software. Experiments were performed three times independently, and data shown are representative of all replicates combined, n ≥ 68 HIEs per condition.

**Dye loading scratch assay.** Lucifer Yellow in Hank's Buffered Salt Solution (HBSS) (1 mg/mL) or Rhodamine 123 in $H_2O$ (1 mg/mL) was diluted 1:100 in FB-Plus and added to MA104 or jHIE cell monolayers. Monolayers were scraped with a 26 *g* needle and incubated for 5 min at room temperature. Monolayers were then washed 3 times with FB-Plus and imaged. Images representative of each experiment were performed in triplicate.

**Immunofluorescence**. MA104 cells and HIEs were fixed using the Cytofix/Cytoperm kit (BD Biosciences) according to manufacturer instructions. Primary antibodies were diluted in 1X Perm/Wash overnight at 4°C. The next day, the cells were washed three times with 1X Perm/Wash solution and then incubated with corresponding secondary antibodies for 1 hr at room temperature. Nuclei were stained with NucBlue Fixed Cell Stain (Life Technologies) for 5 min at room temperature and washed with 1X PBS for imaging. Experiments were performed three times independently.

**Fluorescence focus assay.** Fluorescence focus assays (FFA) were performed as described previously with the following modifications (*13*). After inoculation with RV, MA104 cells were treated with DMSO, 10U/mL apyrase, 10 µM BPTU, 10 µM AR-C 118925XX, 20 µM Bx430, 20 μM 5-BDBD, 300 μM suramin, or 10 μM PPADS. Then, cells and media at 24 hpi were freeze-thawed three times and then with activated with 10 μg/mL Worthington's Trypsin for 30 min at 37 °C prior to use. Ten-fold dilutions were used to inoculate MA104 monolayers in 96-well plates for 1 hr and cells were fixed at 20 hpi with ice-cold methanol for ~20 min at 4°C, then washed 3 times with 1X phosphate buffered solution (PBS). Primary rabbit anti-rotavirus antisera(*12*) was used at 1:100,000 and incubated overnight at 4°C. Monolayers were washed 3x with 1XPBS and then incubated with secondary antibody donkey anti-rabbit AlexaFluor 488 (1:2000) for 1 hr at 37°C and then washed 3x with 1XPBS prior to imaging. FFA experiments were performed three times independently.

**Plaque assay.** Plaque assays were performed as described previously with the following modifications (*59*). Briefly, MA104 cells were seeded and grown to confluency in 6 well plates. Wells were infected at 10-fold dilutions in duplicate for 1 hr and media replaced with an overlay of 1.2% Avicel in serum-free DMEM supplemented with DEAE dextran, and 1μg/mL Worthington's Trypsin, and, for indicated experiments, DMSO vehicle, apyrase, or purinergic receptor blocker drugs (*60*). The cells were incubated at 37°C/5% $CO_2$ for 48-72 hrs before overlay was removed and cells stained with crystal violet to count plaques. Plaque assays of using MA104-GCaMP and MA104-GCaMP-P2Y1ko cells were performed three times independently, and data shown are representative of all replicates combined.

**RNA extraction, reverse transcription, and quantitative PCR.** Total RNA was extracted from HIE 96-well monolayers or MA104 cells grown to confluency in a 6-well plate using TRIzol reagent (Ambion). In RV-infection experiments, cells were harvested at 24 hpi. Total RNA was treated with Turbo DNase I (Ambion) and cDNA was generated from 250 ng RNA using the SensiFAST cDNA synthesis kit (Bioline). Quantitative PCR was performed using Fast SYBR Green (Life Technologies) with primers (**Table S2**) and using the QuantStudio real time thermocycler (Applied

Biosciences). Target genes were normalized to the housekeeping gene ribosomal subunit 18s and relative expression was calculated using the ddCT method. Experiments were performed three times independently, and data shown are representative of all replicates combined. For *IL-1α*, *COX2*, and *iNOS* experiments, HIE monolayers were Mock or RV (Ito)-infected and then treated with DMSO, 100U/mL apyrase, or 10 μM BPTU and normalized to 18s mRNA transcripts and relative to the Mock-DMSO condition.

**Measurement of serotonin release.** Serotonin secretion by HIEs following stimulation with RV infection and treatment with DMSO, 100U/mL apyrase, 10 μM BPTU, or 300 nM ω-agatoxin was quantified by ELISA (Eagle Biosciences) according to the manufacturer's instructions. A standard curve of known serotonin concentrations was plotted against optical density at 450 nm with a limit of detection of 2.6 ng/mL (Infinite F200Pro, Tecan). Experiments in flat monolayers were performed three times independently and experiments in transwells were performed three times independently and data shown are representative of all replicates combined.

**Statistical analysis.** Biostatistical analyses were performed using GraphPad Prism (version 8.1) software (GraphPad Inc., La Jolla, CA). Outlier data points were removed using the ROUT method (Q = 1%). Statistical comparisons were made using Mann- Whitney test, one-way Analysis of Variance (ANOVA), or the Kruskal-Wallis test and the Tukey, Bonferroni, or Dunn's multiple comparisons test. Differences between the groups were considered significant at $p < 0.05$ (*), and the data are presented as mean ± standard deviation unless otherwise noted. All authors had access to the study data, reviewed, and approved the final manuscript.

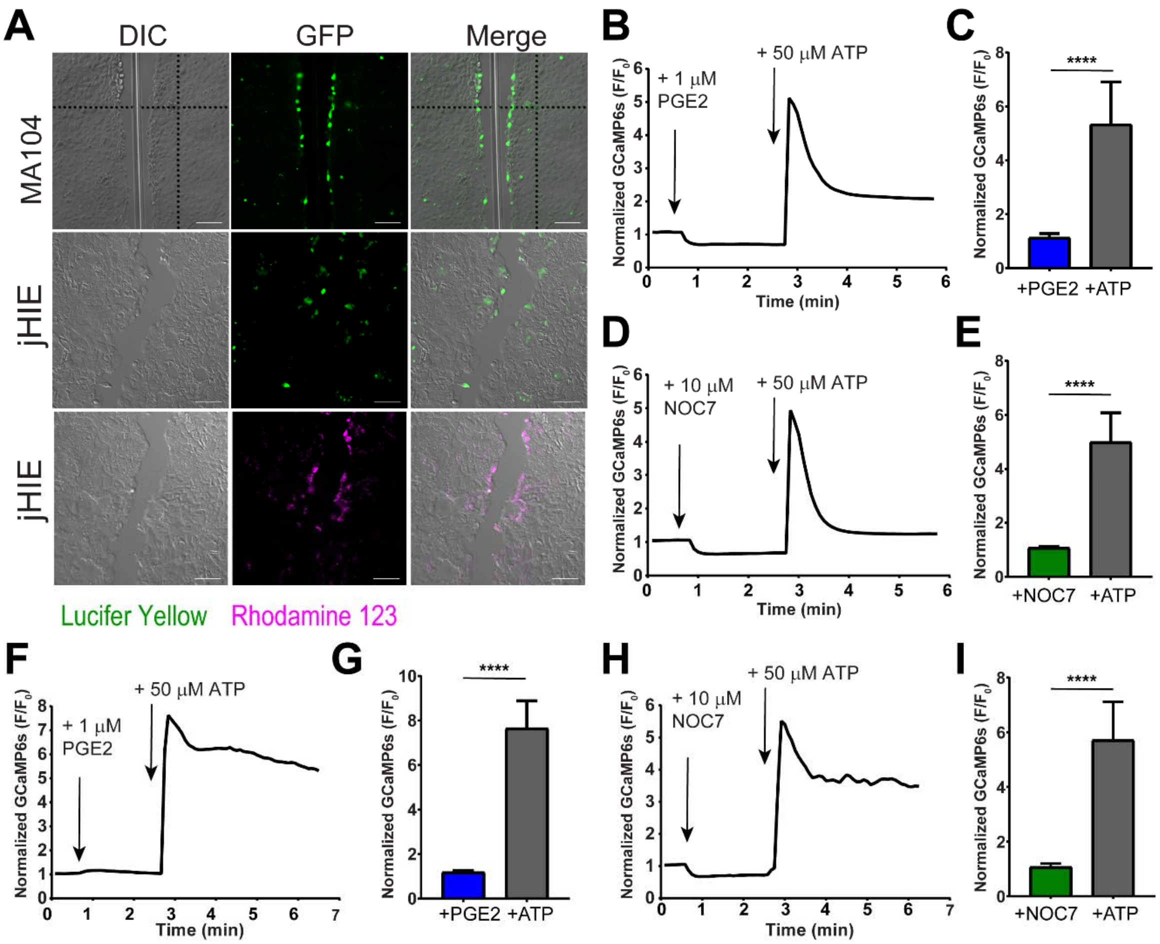


**Figure S1. Rotavirus-induced $Ca^{2+}$ waves do not occur *via* gap junctions, prostaglandin E2, or nitric oxide.**

**(A)** Dye loading scratch assay with Lucifer Yellow and Rhodamine 123 in MA104 cells and jHIE (t J3) monolayers. scale bar = 100 µm. **(B-C)** MA104-GCaMP6s cells treated with 1 µM prostaglandin E2 (PGE2) followed by 50 µM ATP with **(B)** normalized GFP fluorescence average trace and **(C)** maximum normalized GCaMP6s fluorescence increase. **(D-E)** MA104-GCaMP6s cells treated with 10 µM NOC7 followed by 50 µM ATP with **(D)** normalized GFP fluorescence average trace and **(E)** maximum normalized GCaMP6s fluorescence increase. **(F-G)** jHIE-GCaMP6s cells (J2) treated with 1 µM PGE2 followed by 50 µM ATP with **(F)** normalized GFP fluorescence average trace and **(G)** maximum normalized GCaMP6s fluorescence increase. **(H-I)** jHIE-GCaMP6s cells (J2) treated with 10 µM NOC7 followed by 50 µM ATP with **(H)** normalized GFP fluorescence average trace and **(I)** maximum normalized GCaMP6s fluorescence increase. (For **B-I**), n = 30 cells, Mann-Whitney test. Data representative of 4 replicate experiments (****p<0.0001).

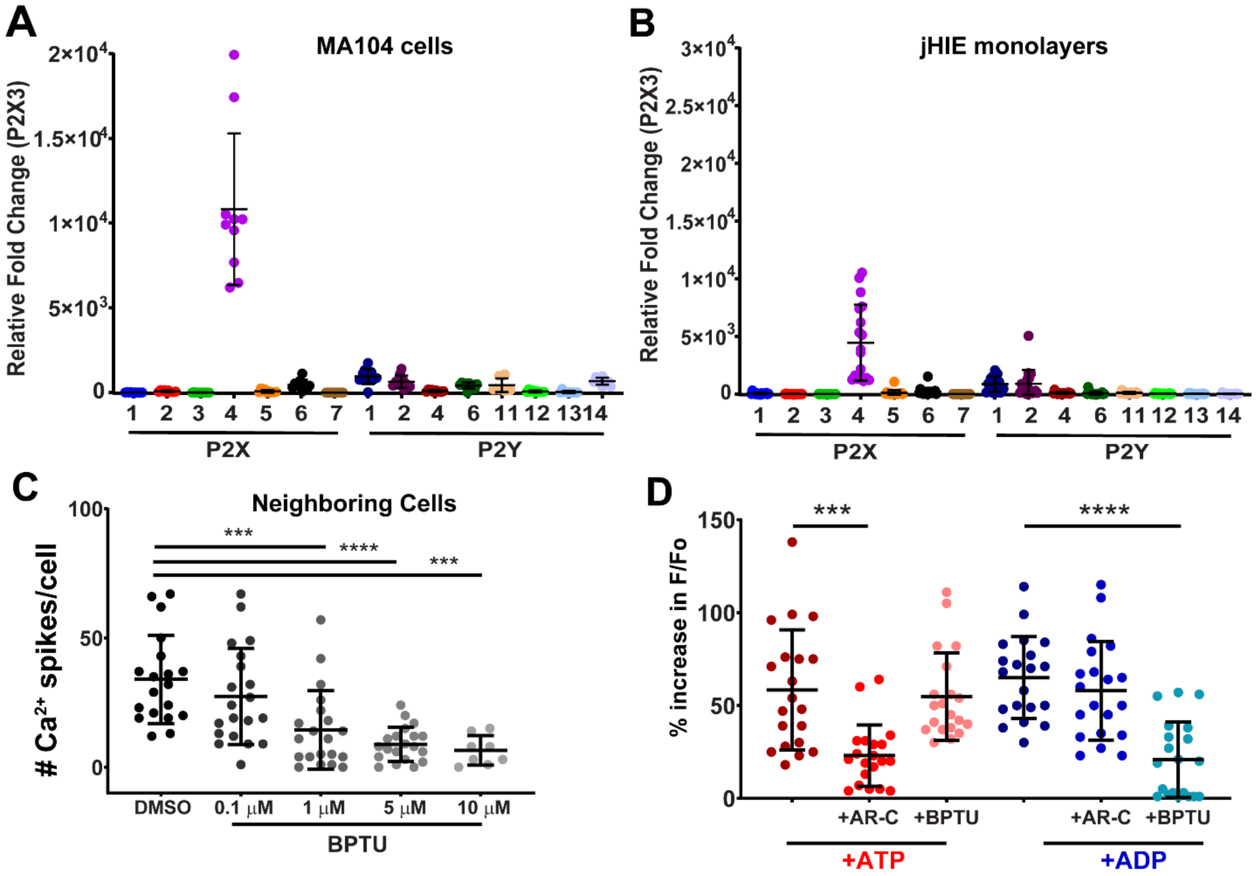


**Figure S2. Purinergic blockers of P2Y1 reduce rotavirus-induced calcium waves.** **(A-B)** qPCR of purinergic receptor mRNA transcripts relative to P2X3 mRNA transcript levels in **(A)** RV (SA114F)-infected MA104 cells or **(B)** RV (Ito)-infected jHIE monolayers. **(C**) $Ca^{2+}$ spikes in NB cells of RV(SA114F)-infected MA104-GCaMP cells treated with DMSO or BPTU. (n= 20 cells, 3 replicates) **(D)** Normalized relative GFP fluorescence of MA104-GCaMP cells incubated with 10 μM AR-C 118925XX (AR-C) or 10 μM BPTU for 3.5 min before addition with 10 nM ADP or 1 μM ATP (n= 20 cells, 3 replicates each). Kruskal-Wallis with Dunn's comparisons test (***p<0.001, ****p<0.0001).

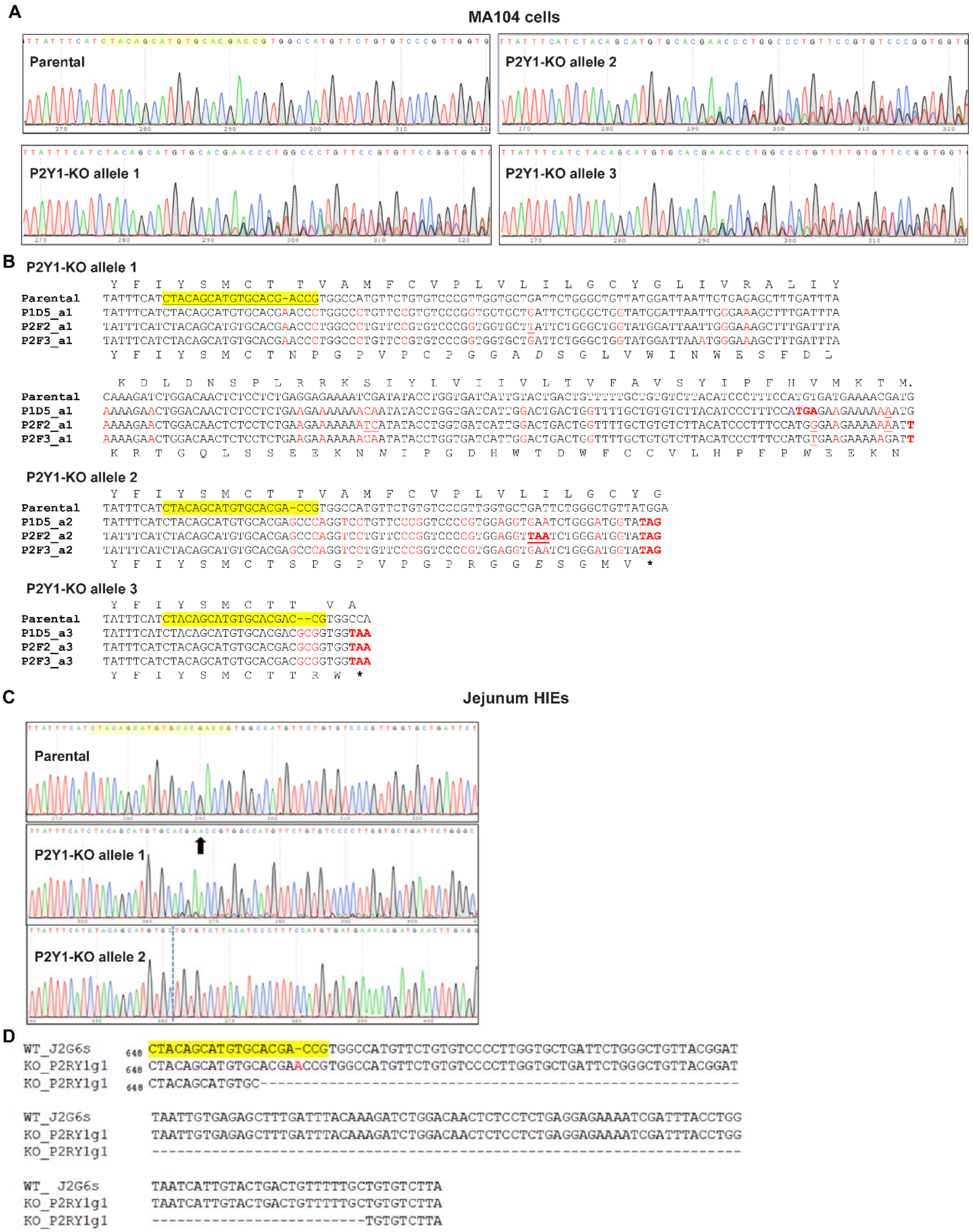


**Figure S3. Genotyping of P2Y1 Knockout Cells.**

**(A)** Sequencing chromatogram and **(B)** genotyping of P2Y1 receptor knockout in MA104-GCaMP-P2Y1ko cells. Parental P2Y1 receptor sequence compared to 3 alleles, mutations in red. **(C)**

Sequence chromatogram and **(D)** genotyping of P2Y1 receptor knockout in jejunum HIE-GCaMP6s-P2Y1ko cells. Parental P2Y1 receptor sequence compared to 2 alleles of the P2Y1 receptor knockouts with an insertion (black arrow) and deletion (dashed line) indicated. The sequence shown is 648 nucleotides relative to the P2Y1 receptor start codon. Small guide RNA sequences highlighted in yellow.

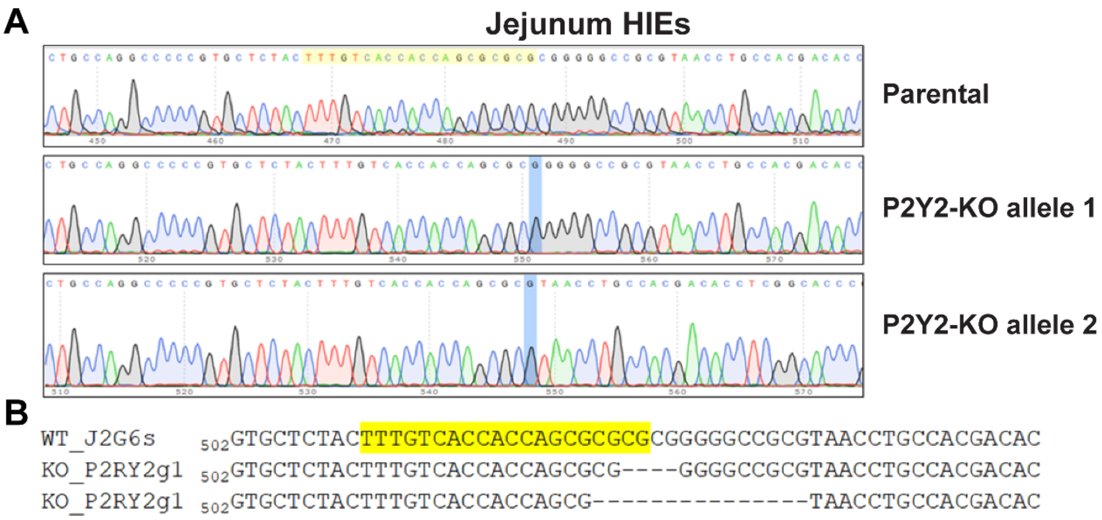


**Figure S4. Genotyping of P2Y2 Knockout HIEs.**
**(A)** Sequence chromatogram and **(B)** genotyping of P2Y2 receptor knockout in jejunum HIE-GCaMP6s-P2Y2ko cells. Parental P2Y2 receptor sequence is compared to 2 alleles from the P2Y2 receptor knockouts with the sites of mutation highlighted in blue. Small guide RNA sequence highlighted in yellow. The sequence shown is 502 nucleotides relative to the P2Y2 receptor start codon.

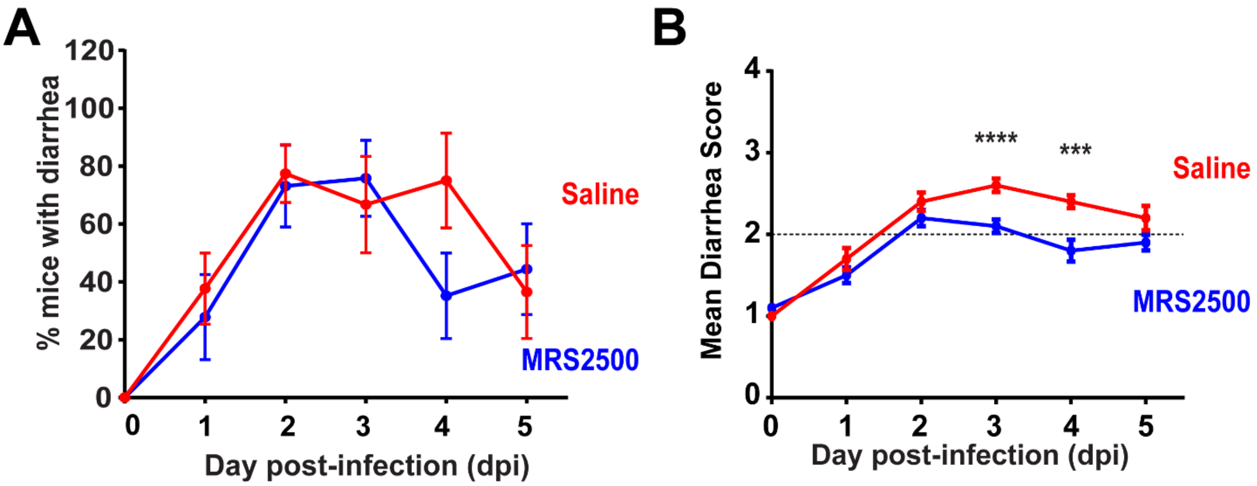


**Figure S5. P2Y1 receptor blocker MRS2500 attenuates rotavirus diarrhea in neonatal mice. (A)** Percentage of C57Bl/6J mouse pups with diarrhea infected with Rhesus RV and vehicle- (8 cages, n = 46 pups) or MRS2500-treated (4 mg/kg) (9 cages, n = 60 pups) and the **(B)** mean diarrhea score. Mann-Whitney test, data presented as mean ± SEM (*p<0.05, **p<0.01).

**Table S1: Genotyping primers**

| Gene Target | Species | Product size | Forward (5') Sequence | Reverse (3') Sequence |
|---|---|---|---|---|
| *P2RY1* | Human | 954 bp | CAGACTGGATCTTCGGGG ATGCC | CCCGCCAAGAAATAGAGAATGGGG |
| *P2RY2* | Human | 1025 bp | CCTGGAATGACACCATCA ATGGC | CCTCTGCATGTCAGTTCTGTCG |

**Table S2: qPCR primer sequences**

| Gene | Species | Forward (5') Sequence | Reverse (3') Sequence | Citation |
|---|---|---|---|---|
| *18S* | Universal | CGCCTTCCTCTTCGAGTATGA | AGATAACGCCCACCTTCTTATT ACG | |
| *IL-1α* | Human | GAATGACGCCCTCAATCAAAGT | TCATCTTGGGCAGTCACATACA | |
| *COX2* | Human | ATCATTCACCAGGCAAATTGC | GGCTTCAGCATAAAGCGTTTG | |
| *iNOS* | Human | CAGCTCCACAAGCTGGCTCG | CAGGATGTCCTGAACGTAGACC TTG | |
| *P2X1* | Human | CGCCTTCCTCTTCGAGTATGA | AGATAACGCCCACCTTCTTATT ACG | |
| *P2X2* | Human | GCCTACGGGATCCGCATT | TGGTGGGAATCAGGCTGAAC | |
| *P2X3* | Human | GCTGGACCATCGGGATCA | GAAAACCCACCCTACAAAGTAGGA | |
| *P2X4* | Human | CCTCTGCTTGCCCAGGTACTC | CCAGGAGATACGTTGTGCTCAA | |
| *P2X5* | Human | CTGCCTGTCGCTGTTCGA | GCAGGCCCACCTTCTTGTT | |
| *P2X6* | Human | AGGCCAGTGTGTGGTGTTCA | TCTCCACTGGGCACCAACTC | |
| *P2X7* | Human | TCTTCGTGATGACAAACTTTCTCAA | GTCCTGCGGGTGGGATACT | |
| *P2Y1* | Human | CGTGCTGGTGTGGCTCATT | GGACCCCGGTACCTGAGTAGA | 61 |
| *P2Y2* | Human | GAACTGACATGCAGAGGATAGAAG AT | GCCGGCGTGGACTCTGT | |
| *P2Y4* | Human | CCGTCCTGTGCCATGACA | TGACCGCCGAGCTGAAGT | |
| *P2Y6* | Human | GCCGGCGACCACATGA | GACCCTGCCTCTGCCATTT | |
| *P2Y11* | Human | CTGGAGCGCTTCCTCTTCAC | GGTAGCGGTTGAGGCTGATG | |
| *P2Y12* | Human | AGGTCCTCTTCCCACTGCTCTA | CATCGCCAGGCCATTTGT | |
| *P2Y13* | Human | GAGACACTCGGATAGTACAGCTGG TA | GCAGGATGCCGGTCAAGA | |
| *P2Y14* | Human | TTCCTTTCAAGATCCTTGGTGACT | GCAGAGACCCTGCACACAAA | |
| *P2X1* | Mouse | CAGTTCTCTGTGATCTCTTATTG | GTCCTCCGCATACTTGAA | |
| *P2X2* | Mouse | TTCACCATCCTCATCAAG | TCAGGTAGTCACTCTTCT | |
| *P2X3* | Mouse | CCCTATTTGTGCTATTAG | TTCTATTCTGTTCTCTTAC | |
| *P2X4* | Mouse | TACGAGCAGGGTCTTTCCGGA | ACAAGACGTGCTCGGGCAAC | |
| *P2X5* | Mouse | CCACACTATTATTTCAAC | CCATTAACTATCACATCA | |
| *P2X6* | Mouse | CTCAAGCTCTATGGAATC | CTTGCCTCTTCATATTTG | |
| *P2X7* | Mouse | GAACCCAAGCCGACGTTGAAGT | GACAGCCTAGACAGGTCGGAGA | |
| *P2Y1* | Mouse | CGCACACAGGTACAGTGGCGT | TTCCGAGTCCCAGTGCCAGAGT | 62 |
| *P2Y2* | Mouse | GTGCGGGGAACCCGGATCAC | AGCCGCCTGGCCATAAGCAC | |
| *P2Y4* | Mouse | GTGTGCACCGCTACATGGGCA | TCAGGCAGAGTCGTGTCATGGCA | |
| *P2Y6* | Mouse | AGGGGACCACTGGCCCTTCG | TACCCAAGCAGCACGGCGAC | |
| *P2Y11* | Mouse | CAGAGTTTGGTCTAGTTTGGCACG A | TGGGAACTTTGGCTGAACCCCT | |
| *P2Y12* | Mouse | GGGCCTCATCGCTTTCGACAGG | TCACGGATGATGGCGTTGCCT | |
| *P2Y13* | Mouse | GGGGCGGAAGTGGCACAAGG | GCGGCTGGACTTCCTCTTGACG | |
| *P2Y14* | Mouse | CAGTTCTCTGTGATCTCTTATTG | GTCCTCCGCATACTTGAA | |

**Supplementary Movie S1. Rotavirus infection induces calcium signaling beyond the infected cell.**
MA104-GCaMP cells were mock- or rotavirus (strain SA114F)-infected and live time-lapse imaging was performed. GCaMP reports cytoplasmic $Ca^{2+}$ as changes in fluorescence intensity (green) and images of RV-antigen positive cells by immunofluorescence (pink) were superimposed on the movies. Note the imbedded timer displays time from the beginning of acquisition not the time post-infection.

**Supplementary Movie S2. Rotavirus-infected cells elicit intercellular calcium waves.**
MA104-GCaMP5G cells were mock-inoculated or rotavirus-infected with the recombinant SA11cl3-mRuby3 reporter virus and live time-lapse imaging was performed. GCaMP5G reports cytoplasmic $Ca^{2+}$ as changes in fluorescence intensity (green) and rotavirus protein synthesis is reported by mRuby3 expression (pink) from the nonstructural protein 3 (NSP3) open-reading frame. Note the imbedded timer displays time from the beginning of acquisition not the time post-infection.

**Supplementary Movie S3. Blocking enterotoxin NSP4 signaling does not reduce intercellular calcium waves in rotavirus infection.**
MA104-GCaMP cells were mock-inoculated or rotavirus-infected with the recombinant SA11cl3-mRuby3 reporter virus and live time-lapse imaging was performed. GCaMP5G reports cytoplasmic $Ca^{2+}$ as changes in fluorescence intensity (green) and rotavirus protein synthesis is reported by mRuby3 expression (pink) from the nonstructural protein 3 (NSP3) open-reading frame. Cells were treated with anti-VP7 M60 MAb, anti-NSP4 MAb 622, or anti-NSP4 antisera 120-147 (Rb Ab) after infection. Note the imbedded timer displays time from the beginning of acquisition not the time post-infection.

**Supplementary Movie S4. Blocking purinergic signaling inhibits intercellular calcium waves in rotavirus infection.**
MA104-GCaMP cells were mock-inoculated or rotavirus-infected with the recombinant SA11cl3-mRuby3 reporter virus and live time-lapse imaging was performed. GCaMP5G reports cytoplasmic $Ca^{2+}$ as changes in fluorescence intensity (green) and rotavirus protein synthesis is reported by mRuby3 expression (pink) from the nonstructural protein 3 (NSP3) open-reading frame. Cells were treated with 10 U/mL apyrase after infection. Note the imbedded timer displays time from the beginning of acquisition not the time post-infection.

**Supplementary Movie S5. Blocking the P2Y1 receptor inhibits intercellular calcium waves in rotavirus infection.**
MA104-GCaMP cells were mock-inoculated or rotavirus-infected with the recombinant SA11cl3-mRuby3 reporter virus and live time-lapse imaging was performed. GCaMP5G reports cytoplasmic $Ca^{2+}$ as changes in fluorescence intensity (green) and rotavirus protein synthesis is reported by mRuby3 expression (pink) from the nonstructural protein 3 (NSP3) open-reading frame. Cells were treated with 10 μM BPTU after infection. Note the imbedded timer displays time from the beginning of acquisition not the time post-infection.

**Supplementary Movie S6. Rotavirus infection induces intercellular calcium waves in human intestinal enteroids.**

Jejunum HIE-GCaMP6s enteroid monolayers were mock- or rotavirus (strain Ito)-infected, and imaged once per minute for ~7-22 hpi. GCaMP6s reports cytoplasmic $Ca^{2+}$ as changes in fluorescence intensity (green). Note the imbedded timer displays time from the beginning of acquisition not the time post-infection.

**Supplementary Movie S7. Blocking purinergic signaling and P2Y1 receptor inhibits intercellular calcium waves in human intestinal enteroids.**

Jejunum HIE-GCaMP6s enteroid monolayers were mock- or rotavirus (strain Ito)-infected, and treated with vehicle (DMSO), 100 U/mL apyrase, or 10 µM BPTU and imaged once per minute for ~7-22 hpi. GCaMP6s reports cytoplasmic $Ca^{2+}$ as changes in fluorescence intensity (green). Note the imbedded timer displays time from the beginning of acquisition not the time post-infection.

**Supplementary Movie S8. CRISPR/Cas9 knockout of the P2Y1 receptor reduces intercellular calcium waves.**

MA104-GCaMP6s or MA104-GCaMP6s-P2Y1ko cells were mock-inoculated or rotavirus-infected with the recombinant SA11cl3-mRuby3 reporter virus and live time-lapse imaging was performed. GCaMP5G reports cytoplasmic $Ca^{2+}$ as changes in fluorescence intensity (green) and rotavirus protein synthesis is reported by mRuby3 expression (pink) from the nonstructural protein 3 (NSP3) open-reading frame. Note the imbedded timer displays time from the beginning of acquisition not the time post-infection.

**Supplementary Movie S9. CRISPR/Cas9 knockout of the P2Y1 receptor inhibits intercellular calcium waves in human intestinal enteroids.**

Jejunum HIE-GCaMP6s, jHIE-GCaMP6s-P2Y1ko, and jHIE-GCaMP6s-P2Y2ko monolayers were mock- or rotavirus (strain Ito)-infected, imaged once per minute for ~7-22 hpi. GCaMP6s reports cytoplasmic $Ca^{2+}$ as changes in fluorescence intensity (green). Note the imbedded timer displays time from the beginning of acquisition not the time post-infection.

**Supplementary Movie S10. Blocking the P2Y1 receptor decreases RV-induced human intestinal enteroid swelling.**

3D jejunum HIE-GCaMP6s enteroids were mock- or rotavirus (strain Ito)-infected, treated with vehicle (DMSO) or 10 µM BPTU, and imaged once per 2-3 min in GFP and differential interference contrast on a widefield epifluorescence microscope for ~3-21 hpi. GCaMP6s reports cytoplasmic $Ca^{2+}$ as changes in fluorescence intensity (green). Note the imbedded timer displays time from the beginning of acquisition not the time post-infection.
